# NPC1-mTORC1 signaling Couples Cholesterol Sensing to Organelle Homeostasis and is a Targetable Pathway in Niemann-Pick type C

**DOI:** 10.1101/2020.08.02.233254

**Authors:** Oliver B. Davis, Hijai R. Shin, Chun-Yan Lim, Emma Y. Wu, Matthew Kukurugya, Claire F. Maher, Rushika M. Perera, M. Paulina Ordonez, Roberto Zoncu

## Abstract

Lysosomes promote cellular homeostasis through macromolecular hydrolysis within their lumen and metabolic signaling by the mTORC1 kinase on their limiting membranes. Both hydrolytic and signaling functions require precise regulation of lysosomal cholesterol content. In Niemann-Pick type C (NPC), loss of the cholesterol exporter, NPC1, causes cholesterol accumulation within lysosomes, leading to mTORC1 hyperactivation, disrupted mitochondrial function and neurodegeneration. The compositional and functional alterations in NPC lysosomes, and how aberrant cholesterol-mTORC1 signaling contributes to organelle pathogenesis are not understood. Through proteomic profiling of NPC lysosomes, we find pronounced proteolytic impairment compounded with hydrolase depletion and enhanced membrane damage. Genetic and pharmacologic mTORC1 inhibition restores lysosomal proteolysis without correcting cholesterol storage, implicating aberrant mTORC1 as a pathogenic driver downstream of cholesterol accumulation. Consistently, mTORC1 inhibition ameliorates mitochondrial dysfunction in a neuronal model of NPC. Thus, cholesterol-mTORC1 signaling controls organelle homeostasis and is a targetable pathway in NPC.

## INTRODUCTION

Lysosomes are degradative organelles that play key roles in macromolecular turnover and in recycling of cellular building blocks including amino acids, lipids and nucleotides. Through their participation in autophagy, lysosomes also help detoxify potentially harmful cellular components, such as damaged mitochondria. In addition to their recycling and degradative roles, lysosomes have been recently recognized as signaling compartments that support the activation and regulation of the master growth regulator, mechanistic Target of Rapamycin Complex 1 (mTORC1) (Liu and Sabatini, 2020; Perera and Zoncu, 2016).

The degradative and signaling roles of lysosomes are highly integrated. Intracellular nutrients drive mTORC1 translocation from the cytosol to the lysosomal limiting membrane, where growth factor signals relayed by the phosphatidylinositol 3-kinase (PI3K)-AKT pathway trigger the kinase function of mTORC1 (Castellano et al., 2017; Menon et al., 2014; Sancak et al., 2010; Wyant et al., 2017; Zoncu et al., 2011). Moreover, upon becoming activated at the lysosome by converging nutrient- and growth factor-mediated inputs, mTORC1 strongly suppresses initiation of autophagy, steering the cell toward net mass accumulation (Düvel et al., 2010; Kim et al., 2011; Puente et al., 2016; Settembre et al., 2012). The tight integration of the recycling and signaling functions of the lysosome suggests that, in diseases driven by lysosomal dysfunction, aberrant regulation of both processes may synergize to disrupt cellular homeostasis (Perera and Zoncu, 2016). However, the respective roles of aberrant lysosomal recycling and signaling, and their interplay in driving disease pathogenesis, remain largely unexplored.

Niemann-Pick type C (NPC) is one of a family of approximately 60 diseases known as lysosomal storage disorders (LSDs), in which genetic inactivation of lysosomal hydrolases or transporters triggers massive and pathogenic accumulation of their respective substrates within the lysosome (Ballabio and Gieselmann, 2009; Platt et al., 2018). NPC is triggered by inactivating mutations in NPC1, a polytopic transmembrane cholesterol transporter located on the lysosomal limiting membrane (Gong et al., 2016; Kwon et al., 2009; Li et al., 2016; Winkler et al., 2019). In conjunction with NPC2, a cholesterol-binding protein of the lysosomal lumen, NPC1 exports cholesterol released from Low-Density Lipoprotein (LDL) to acceptor compartments such as the endoplasmic reticulum (ER), the Golgi and the plasma membrane (Feltes et al., 2020; Infante and Radhakrishnan, 2017; Infante et al., 2008; Pfeffer, 2019).

A large number of heritable mutations in the NPC1 gene lead to an unstable NPC1 protein, which fails to fold and is degraded in the ER (Schultz et al., 2018). In cells lacking NPC1, cholesterol accumulates massively within the lysosomal lumen, as well as on its limiting membrane. Cholesterol storage results in enlarged lysosomes that exhibit morphological, trafficking and functional defects. Moreover, the primary lysosomal phenotype is accompanied by dysfunction in other cellular compartments, including autophagosomes (Elrick et al., 2012; Ordonez et al., 2012; Sarkar et al., 2013), mitochondria (Kennedy et al., 2014; Ordonez, 2012; Yambire et al., 2019a; Yu et al., 2005) and peroxisomes (Schedin et al., 1997).

Mechanistically, how cholesterol accumulation caused by loss of NPC1 leads to lysosomal dysfunction remains poorly understood. There is evidence of defective degradation of autophagosomal cargo (Elrick et al., 2012; Ordonez et al., 2012), as well as defective trafficking of autophagic vesicles to lysosomes (Sarkar et al., 2013). However, the mechanistic basis for impaired lysosomal catabolism remains to be fully elucidated. Moreover, the molecular processes that connect the primary lysosomal dysfunction to the impairment of other organelle populations remain largely mysterious. It was recently reported that a faulty transcriptional circuit triggered by lipid storage compromises mitochondrial biogenesis in NPC (Yambire et al., 2019a). However, it is likely that additional lysosome-based processes may be involved in the loss of mitochondrial homeostasis.

Limiting our understanding of the pathogenic processes that drive NPC is the lack of a comprehensive view of the structural and functional alterations that occur in each membrane compartment in NPC cells. The recent development of techniques for rapid immunoisolation and systematic mass spectrometry-based profiling of intact organelles presents with an opportunity to address this critical point (Abu-Remaileh et al., 2017; Castellano et al., 2017; Sleat et al., 2013; Wyant et al., 2017; Zoncu et al., 2011).

An especially important question in understanding NPC pathogenesis is whether disruption of a common signaling pathway may underlie the multi-organellar loss of function of NPC1-defective cells. Cholesterol was recently identified as a nutrient input that regulates mTORC1 and promotes activation of its downstream biosynthetic programs as well as suppression of autophagy. Cholesterol stimulates mTORC1 activity via a protein complex also involved in amino acid-dependent mTORC1 activation and composed of the heterodimeric Rag guanosine triphosphatases (GTPases), their membrane-anchored interactor Ragulator complex, and the lysosomal amino acid permease SLC38A9 (Castellano et al., 2017; Sancak et al., 2010; Wang et al., 2015; Wyant et al., 2017). This complex responds to both extracellular cholesterol, carried by low-density lipoprotein (LDL) and to intracellular cholesterol transferred across ER-lysosome contacts by a tag team of oxysterol binding protein (OSBP) and its ER anchors, VAPA and B (Castellano et al., 2017; Dong et al., 2016; Lim et al., 2019; Mesmin et al., 2013).

In contrast to these positive activators, NPC1 appears to antagonize cholesterol-dependent mTORC1 signaling. In NPC1-defective cells, mTORC1 signaling is elevated compared to wild-type cells and is immune to inhibition by cholesterol-depleting agents. mTORC1 dysregulation can be readily explained by the massive accumulation of lysosomal cholesterol resulting from loss of NPC1-dependent export. However, NPC1 could also play a cholesterol effector role, where it could directly regulate the mTORC1-scaffolding complex via physical interaction (Castellano et al., 2017).

mTORC1 signaling has profound effects on organelle composition and function. mTORC1 negatively regulates the basic helix-loop-helix MiT-TFE transcription factors, which are master regulators of lysosomal biogenesis and autophagy (Martina et al., 2012; Roczniak-Ferguson et al., 2012; Sardiello et al., 2009; Settembre et al., 2012). mTORC1 drives mitochondrial biogenesis and function via both transcriptional and translational mechanisms (Bentzinger et al., 2008; Cunningham et al., 2007; Morita et al., 2013), and stimulates mitochondrial metabolic programs that support cell growth and proliferation (Ben-Sahra et al., 2016; Csibi et al., 2013). Thus, dysregulation of mTORC1 signaling due to loss of NPC1 could affect organelle homeostasis via multiple mechanisms.

To shed light into the relationship between dysregulated mTORC1 signaling and organelle homeostasis in NPC, we carried out mass spectrometry-based profiling of immunopurified lysosomes combined with functional assays for mTORC1 signaling and lysosome function in both engineered cell lines and in iPSC-derived neuronal cultures lacking NPC1. We uncover a more profound disruption of lysosomal composition and function, mitochondrial homeostasis and cholesterol-mTORC1 signaling associated with NPC than previously anticipated. Importantly, pharmacological suppression of mTORC1 signaling was able to correct several aspects of organelle function independent of the primary cholesterol storage defect, thus implicating dysregulated mTORC1 signaling as a likely pathogenic driver in Niemann-Pick type C.

## RESULTS

### 1. NPC1-null lysosomes display extensive proteolytic defects

To investigate the molecular basis for lysosomal dysfunction in NPC, we conducted lysosome immunoprecipitation (lyso-IP) followed by label-free proteomics-based profiling from HEK-293T cells that either have intact NPC1 function or in which the NPC1 gene was targeted using CRISPR/Cas9, resulting in near-complete (∼93%) loss of NPC1 protein levels (Table S1) and establishment of an NPC-like phenotype characterized by massive cholesterol accumulation in lysosomes (Figure S1A) (Abu-Remaileh et al., 2017; Castellano et al., 2017; Lim et al., 2019; Wyant et al., 2017; Zoncu et al., 2011). To identify cargo proteins being actively degraded, we incorporated additional control samples treated with the broad-spectrum hydrolase inhibitors, leupeptin and pepstatin (Figure S1B and S1C). In lysosomes purified from control (‘sgNT’) cells, the abundance of resident lysosomal proteins (hydrolases, permeases, signaling components) remained unchanged. In contrast, lysosomal cargos, were enriched 2-fold or more in the leupeptin:pepstatin-treated samples (Figure 1A). These included autophagic adaptors, likely delivered to the lysosome via fusion with autophagosomes (Khaminets et al., 2016). Unlike lysosomes isolated from control cells, NPC1-defective lysosomes showed reduced enrichment of numerous *bona fide* lysosomal cargos in the leupeptin:pepstatin condition (Figure 1B), as most lysosomal cargo were already accumulated within NPC1-null lysosomes in the vehicle treated condition (Fig 1C).

**Figure 1.**
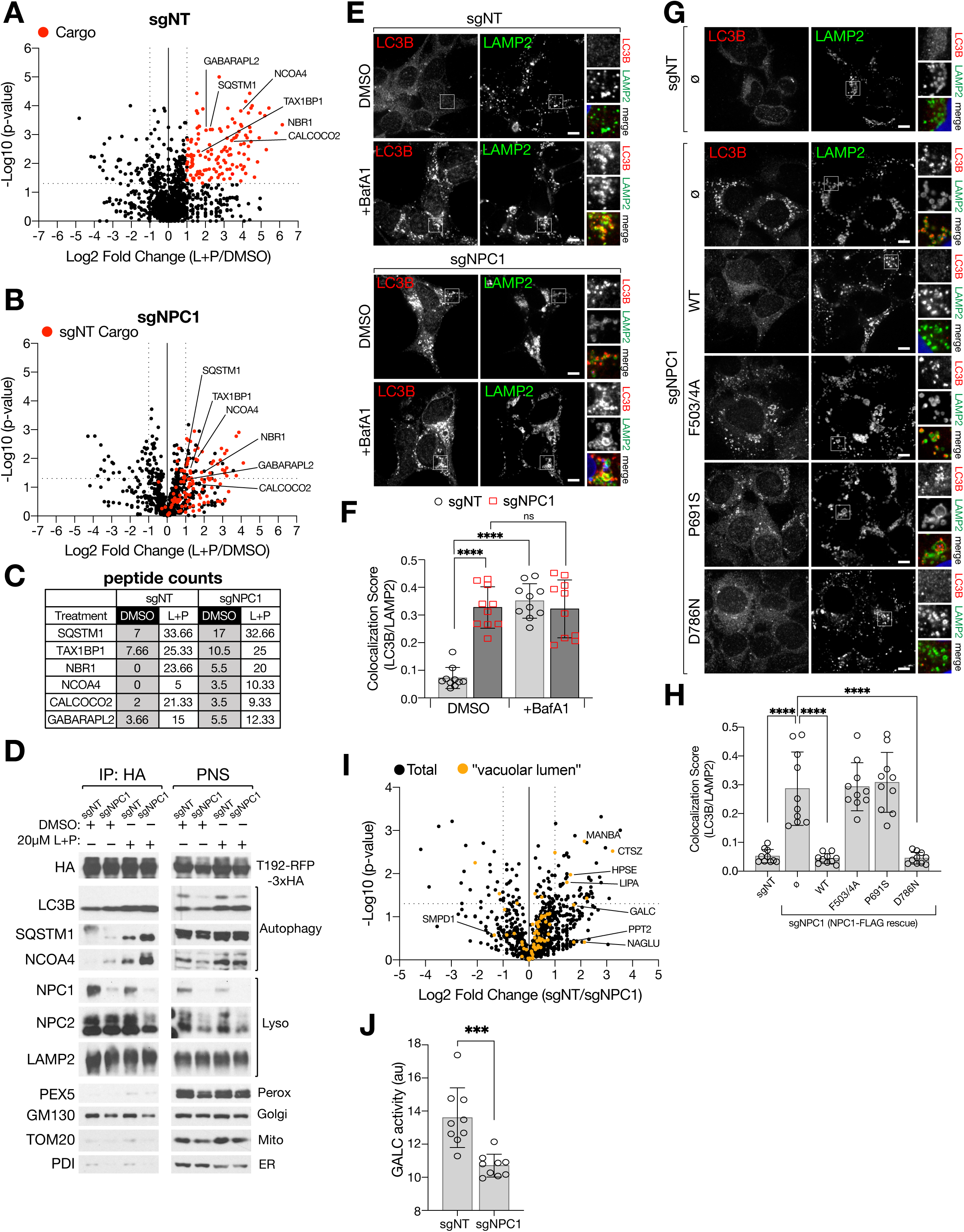
NPC1-deficient lysosomes have reduced degradative capacity relative to sgNT lysosomes. (A-C) Proteomic analysis of Lyso-IP samples from sgNT (NT = “non-targeting” control) and sgNPC1 293Ts. Volcano plots (-log10 p-value vs. log2 of the ratio of leupeptin+pepstatin/DMSO) for sgNT (A) and sgNPC1 (B) 293Ts. Proteins with statistically significant (p-value ≥ 0.5, two-tailed unpaired t-test) fold change L+P/DMSO > 2 (“sgNT cargo”) in (A) are displayed as red circles. The sgNT cargo proteins identified in (A) are also displayed as red circles in (B). (C) Average peptide counts (raw) for selected autophagy-related proteins. (D) Immunoblots of Lyso-IP samples and corresponding post-nuclear supernatant (PNS) from sgNT or sgNPC1 293Ts (expressing TMEM192-RFP-3xHA) treated with 20µM leupeptin and pepstatin for 24h. (E and F) sgNT and sgNPC1 293Ts were treated with 500nM bafilomycin A1for 4h before being fixed and stained with antibodies targeting LC3B and LAMP2. (E) Representative confocal micrographs for each sample. (F) Quantification of the co-localization between LC3B and LAMP2; ****P < 0.0001, ns = not significant, ANOVA with Tukey’s multiple comparisons test. (G and H) sgNT and sgNPC1 293Ts expressing the indicated NPC1-FLAG cDNA were fixed and stained with antibodies targeting LC3B and LAMP2. (I) Representative confocal micrographs for each sample. (J) Quantification of the co-localization between LC3B and LAMP2; ****P(adjusted) < 0.0001, ANOVA with Dunnett’s multiple comparisons test. (I) Volcano plot of Lyso-IP proteomic data (from A-C) for the ratio of untreated (DMSO) sgNT/sgNPC1 LAMP1-normalized peptide counts. Proteins that are classified by the GO term “vacuolar lumen” (GO:0005775) are depicted as orange circles. Statistical analysis was performed using two-tailed unpaired t-test. (J) Relative galactosylceramidase (GALC) activity in sgNT or sgNPC1 293Ts from cells labeled with a GALC activity probe (GalGreen) and LysoTracker Red. GalGreen fluorescence is normalized by total lysosomal content, as determined by LysoTracker Red fluorescence; ***P = 0.0004, two-tailed unpaired t-test. Scale bars in all images are 10µm.

To validate the proteomic data, we conducted direct immunoblotting of lysosomal immunoprecipitates from both control and NPC1-defective HEK293T cells. We found that the steady-state levels of several autophagic adaptor proteins, including LC3, p62/SQSTM1, TAXBP1 and NCOA4 were significantly elevated in NPC1-null lysosomes compared to control lysosomes (Figure 1D, S1D). Blocking lysosomal proteolysis with leupeptin/pepstatin or with the vacuolar H^+^ATPase (v-ATPase) inhibitor, bafilomycin A1 (BafA1) caused further buildup of autophagic substrates in both genetic backgrounds, however this buildup was significantly reduced in NPC1-null lysosomes (Figure 1D, S1D). Immunoblotting of lysosomal immunoprecipitates from NPC1-knock out mouse embryonic fibroblasts (MEFs) also confirmed increased amounts of undigested autophagic substrates (Figure S1E).

We further confirmed accumulation of undigested autophagic material upon loss of NPC1 via immunofluorescence staining for endogenous LC3B and LAMP2, which showed strong LC3B signal within the lumen of LAMP2-positive vesicles in NPC1-depleted but not control cells (Figure 1E-1F, and S1F-S1G). Consistent with defective proteolysis, BafA1 treatment caused a 5-fold increase of LC3B signal in control lysosomes, but no significant change in NPC1-defective lysosomes (Figure 1E-1F).

The proteolytic defects characteristic of NPC1-null lysosomes are a direct consequence of ablated cholesterol-exporting activity. Reconstituting NPC1-defective cells with either wild-type NPC1 or with a mutant (D786N) that is predicted to have enhanced transport activity based on homology to the ER-resident cholesterol sensor SCAP (Gao et al., 2017; Yabe et al., 2002) fully rescued the LC3B accumulation defect (Figure 1G and 1H). In contrast expression of several NPC1 mutant variants that either fail to export cholesterol due to impaired binding to NPC2 (F503/504A) or have impaired cholesterol binding to the putative sterol-sensing domain (SSD: P691S) (Gong et al., 2016; Li et al., 2016; Millard et al., 2005) failed to clear LC3B-positive material from the lysosomal lumen (Figure 1G and 1H). F503/504A and SSD mutants were correctly targeted to the lysosome membrane (Figure S2A) and, as expected, failed to correct lysosomal cholesterol buildup, as shown by unchanged staining with filipin (Figure S2B) and with the recombinant sterol probe, mCherry-D4H*, which binds to cholesterol on the limiting membrane of the lysosome (Lim et al., 2019; Maekawa and Fairn, 2015) (Figure S2C and S2D).

To dissect the molecular basis for the pronounced proteolysis defect of NPC1-null lysosomes, we quantified the abundance of resident lysosomal proteins. We found that several lumenal hydrolases were decreased or undetectable in the NPC-null compared to control lysosomal samples. These included the Cathepsin Z protease (CTSZ), the acid lipase (LIPA), which de-esterifies LDL-derived cholesterol, and several enzymes involved in degradation of glycans and glycosphingolipids such as galactosylceramidase (GALC) and N-acetyl-alpha-glucosaminidase (NAGLU) (Figure 1I). A fluorescence-based reporter assay for the activity of GALC showed a marked reduction in NPC1-depleted cells compared to control cells (Figure 1J). Taken together, these data indicate that reduced abundance of resident lysosomal hydrolases may lead to impaired degradative capacity of NPC lysosomes and defective autophagic cargo degradation in NPC1 null cells.

### 2. NPC1-null lysosomes display increased susceptibility to membrane damage

Recently, it has emerged that the lysosomal limiting membrane is susceptible to damage resulting from undigested/undigestible substrates that accumulate within the lumen (Jia et al., 2020; Maejima et al., 2013; Radulovic et al., 2018; Skowyra et al., 2018). Thus, we considered possibility that defective lumenal proteolysis upon loss of NPC1 function might compromise the integrity of the lysosomal membrane. An early detection system provided by the Endosomal Sorting Complex Required for Transport (ESCRT) III proteins detects ‘microtears’ in the lysosomal membrane and repairs them via a membrane scission process that is thought to topologically resemble ESCRT III-mediated intraluminal vesicle budding during endosomal maturation (Nguyen et al., 2020; Radulovic et al., 2018; Schöneberg et al., 2017; Skowyra et al., 2018). Accordingly, our proteomics analysis found enrichment of the ESCRT III-interacting protein ALIX (also known as PDCD6IP), along with ESCRT III components CHMP1A and IST1 in NPC1-defective over wild-type lysosomes (Table S1) (Figure 2A). Together with the ESCRT I protein, TSG101, ALIX provides a recruiting platform for the assembly of coiled filaments of ESCRT III subunits, including CHMP1A and IST1, to sites of damage on lysosomal membranes (Radulovic et al., 2018; Skowyra et al., 2018).

**Figure 2.**
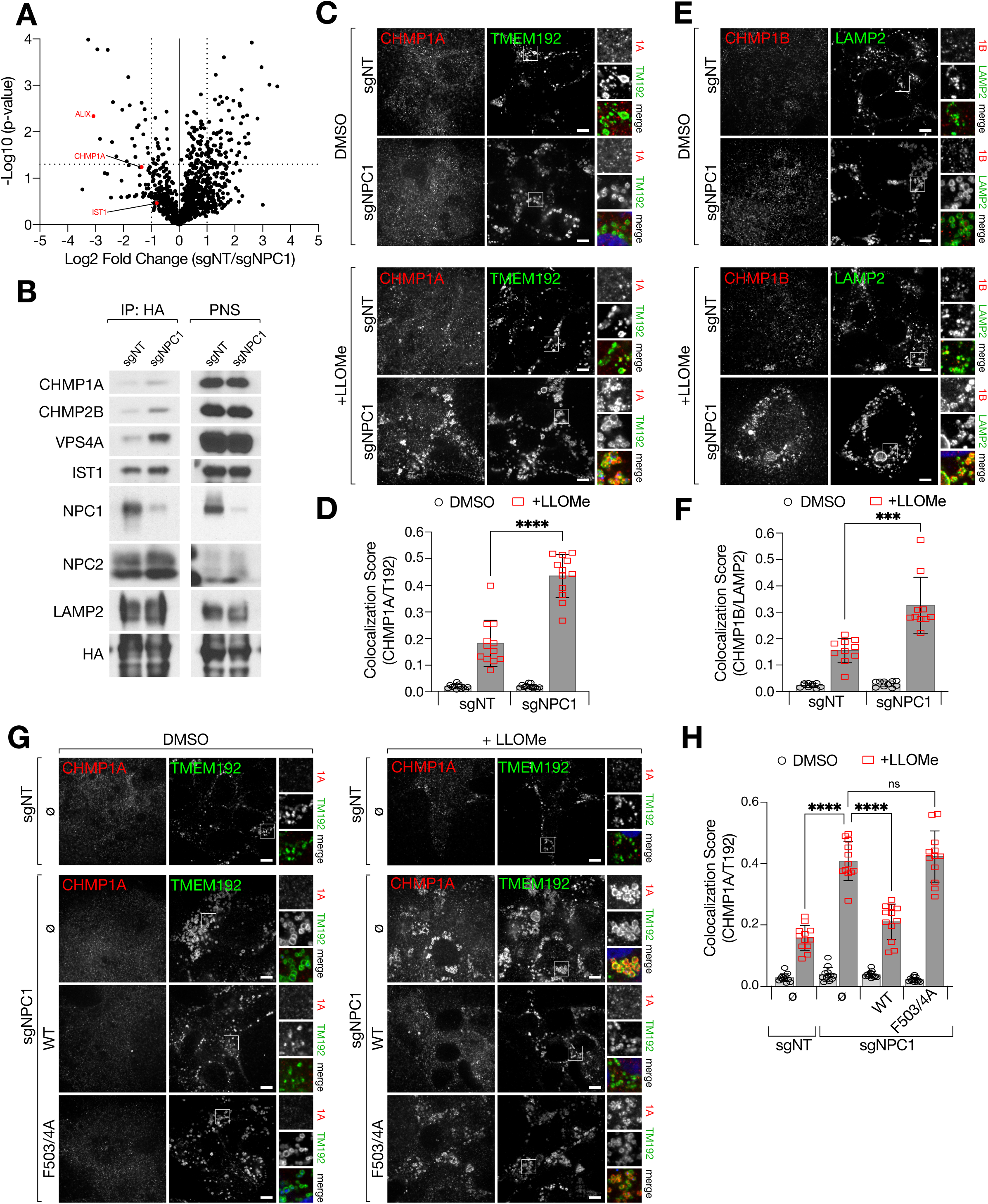
NPC1-deficient lysosomes display an increased propensity for membrane damage. (A) Volcano plot of Lyso-IP proteomic data (from 1A-C) for the ratio of untreated (DMSO) sgNT/sgNPC1 LAMP1-normalized peptide counts. Selected ESCRT proteins are depicted as magenta circles. Statistical analysis was performed using two-tailed unpaired t-test. (B) Immunoblots of Lyso-IP samples from sgNT or sgNPC1 293Ts expressing TMEM192-RFP-3xHA. (C-F) sgNT or sgNPC1 293Ts were treated with 1mM LLOMe or vehicle (DMSO) for 10m before being fixed and stained with the indicated antibodies. Representative confocal micrographs for cells stained with CHMP1A and TMEM192 (C) or CHMP1B and LAMP2 (E) Quantification of the co-localization between CHMP1A and TMEM192 (D) or CHMP1B and LAMP2 (F); ****P < 0.0001, ***P = 0.0005, two-tailed unpaired t-test with Welch’s correction. (G and H) sgNT and sgNPC1 293Ts expressing the indicated NPC1-FLAG cDNA were fixed and stained with antibodies targeting CHMP1A and TMEM192. (G) Representative confocal micrographs for each sample. (H) Quantification of the co-localization between CHMP1A and TMEM192; ****P(adjusted) < 0.0001, ANOVA with Dunnett’s multiple comparisons test. Scale bars in all images are 10µm.

Consistent with the proteomic data, immunoblotting of lysosomal immunoprecipitates from NPC1-defective HEK-293Ts confirmed increased lysosomal accumulation of several ESCRT III components, including CHMP1A, CHMP2B and IST1, as well as the AAA+ ATPase, VPS4A, which mediates ESCRT III polymer remodeling and disassembly (Chiaruttini et al., 2015; Schöneberg et al., 2017) (Figure 2B). In contrast to the clear proteomics and immunoblotting results, ESCRT III accumulation in NPC1-defective cells was not readily visible by double immunofluorescence staining for lysosomal markers and CHMP1A or CHMP1B, possibly due to limitations of the available antibodies. However, enhanced ESCRT III accumulation became evident upon induction of lysosomal damage using the membrane destabilizing agent L-leucyl-L-leucine methyl ester (LLOMe), both in NPC1-defective HEK293Ts and in NPC1-null MEFs (Figure 2C-2F and Figure S3A-S3D). Given that lysosomal pH is not significantly compromised by loss of NPC1 (Elrick et al., 2012), the enrichment of ESCRT III components is likely not reflective of unrecoverable membrane permeabilization, but rather of higher propensity to damage events that are compensated, at least in part, by local ESCRT III polymerization.

As was the case for defective proteolysis, the higher propensity of NPC lysosomes to membrane damage was a direct consequence of the loss of NPC1-dependent cholesterol export. Reconstituting NPC1-defective HEK293T cells and MEFs with wild-type NPC1 suppressed LLOMe-induced lysosomal CHMP1A and CHMP1B accumulation, whereas the transport-defective F503/4A mutant failed to do so (Figure 2G-2H and Figure S3F-S3G).

Thus, our lysosomal proteomics analysis reveals that increased propensity to membrane damage accompanies defective proteolysis and loss of hydrolases following loss of NPC1-dependent cholesterol export. These defects are causally linked to cholesterol accumulation and, quite conceivably, to one another, as accumulation of undigested substrates could lead to membrane rupture through mechanical stress.

### 3. NPC1 regulates mTORC1 via its cholesterol-exporting function

To begin to establish a mechanistic link between the lysosomal defects described above and faulty mTORC1 regulation, we tested the ability of several sterol transport-defective NPC1 mutants to restore mTORC1 regulation by cholesterol. Reconstituting NPC1-depleted cells with wild-type NPC1 rescued the correct regulation of mTORC1 signaling upon cholesterol depletion via MCD, followed by repletion with either free cholesterol or LDL (Figure 3A and S4A). Unlike wild-type, the NPC2-binding defective (F503/504A) and the sterol-sensing domain (P691S) mutants of NPC1 (expressed at nearly identical levels to the wild-type protein) were unable to restore mTORC1 regulation by cholesterol depletion-refeed (Figure 3A and S4A). Another transport-incompetent mutant, caused by impaired cholesterol binding to the N-terminal lumenal domain (P202/3A) (Kwon et al. 2009), also failed to restore mTORC1 sensitivity to cholesterol depletion (Figure S2A-S2B and Figure S4B). Conversely, the transport-competent D786N mutant fully restored cholesterol- or LDL-dependent mTORC1 regulation (Figure 3A and S4A). In agreement with the signaling results, the transport defective NPC1 isoforms failed to re-establish regulation of mTORC1 localization to LAMP2-positive lysosomes in response to changes in cellular cholesterol levels (Figure 3B and 3C).

**Figure 3.**
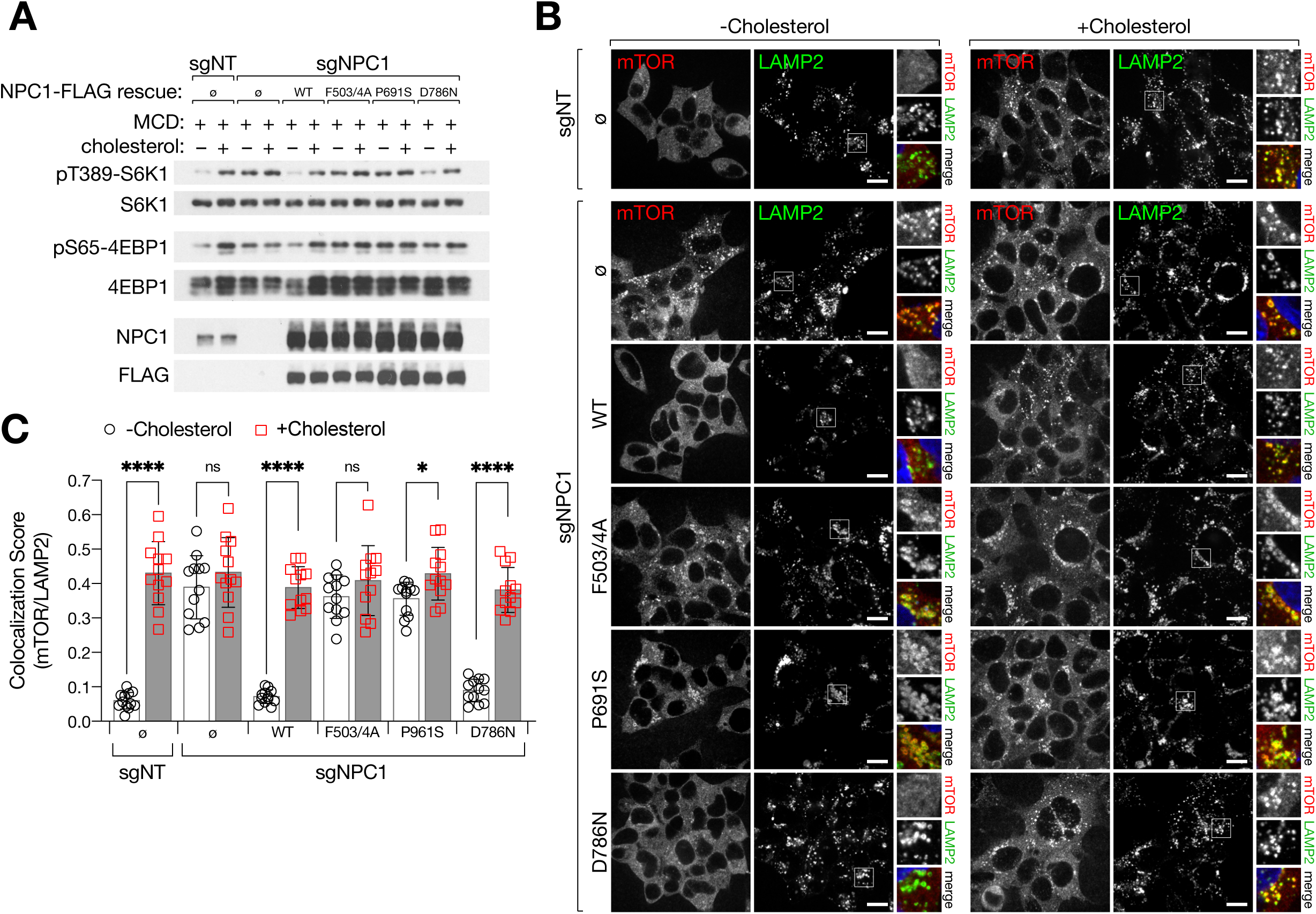
Cholesterol transport by NPC1 controls mTORC1 activity in response to lysosomal cholesterol. (A) Immunoblots from sgNT and sgNPC1 293Ts expressing the indicated NPC1-FLAG cDNA. Cells were depleted of sterols using methyl-β-cyclodextrin (MCD, 0.75% w/v) for 2h, followed by re-feeding for 2h with 50µM cholesterol in complex with 0.1% MCD, as indicated. (B and C) Cells were starved for and re-fed with cholesterol as in (A) before being fixed and stained with antibodies directed against mTOR and LAMP2. (B) Representative confocal micrographs for each sample. (C) Quantification of the co-localization between mTOR and LAMP2; ****P < 0.0001, *P = 0.0121, ns = not significant, two-tailed unpaired t-test with Welch’s correction. Scale bars in all images are 20µm.

Together, these data argue against a sterol-sensing role of NPC1 in the mTORC1 pathway, and strongly suggest that the cholesterol exporting function of NPC1 is necessary and sufficient for mTORC1 regulation.

### 4. mTORC1 inhibition restores integrity and proteolytic function of NPC lysosomes downstream of cholesterol storage

We next investigated the cellular effects of aberrant mTORC1 regulation in NPC. Due to its ability to stimulate lipogenic programs (Düvel et al., 2010; Li et al., 2010; Peterson et al., 2011), conceivably mTORC1 could contribute to lysosomal cholesterol buildup in NPC1-null cells. However, treating NPC cells with the ATP-competitive mTOR inhibitor, Torin1 (Thoreen et al., 2009), did not significantly reduce cholesterol accumulation in the lysosomal lumen or limiting membrane, as indicated by unchanged staining with filipin and mCherry-D4H*, respectively (Figure 4A and 4B). Thus, mTORC1 dysregulation occurs downstream of cholesterol storage and does not appear to contribute significantly to its establishment.

**Figure 4.**
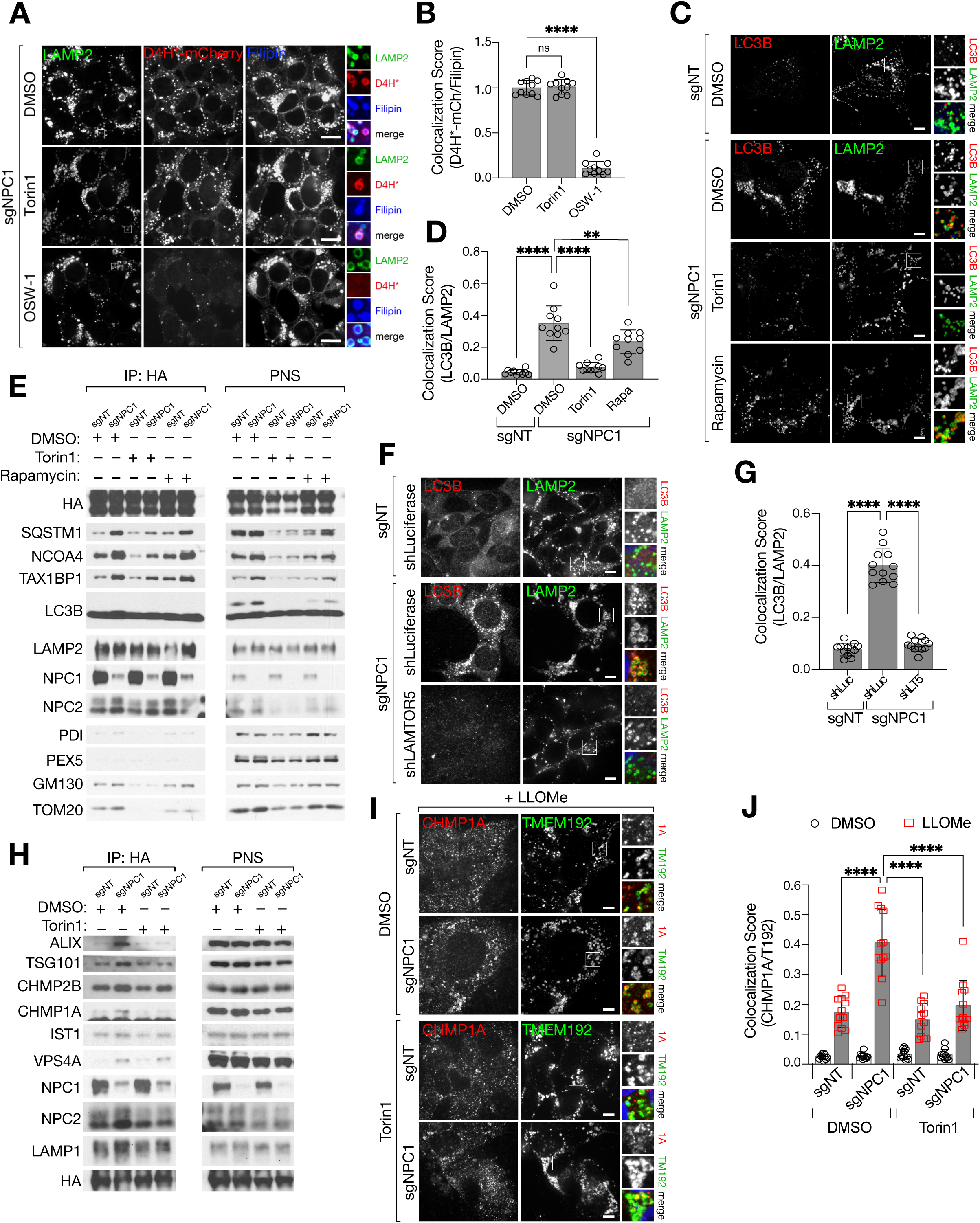
Inhibition of mTORC1 activity alleviates lysosomal pathologies associated with loss of NPC1. (A and B) sgNPC1 293Ts were treated with Torin1 (250nM, 24h), OSW-1 (10nM, 8h), or vehicle (DMSO) before being fixed and semi-permeabilized with a liquid N2 pulse, followed by cholesterol labeling with D4H*-mCherry and filipin, and staining with antibodies directed against LAMP2. (A) Representative confocal micrographs for each sample. Scale bars are 20µm. (B) Quantification of the co-localization of D4H*-mCherry and filipin-positive structures; ****P(adjusted) < 0.0001, ANOVA with Dunnett’s multiple comparisons test. (C and D) sgNT or sgNPC1 293Ts were treated with Torin1 (250nM, 24h), rapamycin (100nM, 24h), or vehicle (DMSO) as indicated before being fixed and stained for antibodies directed against LC3B and LAMP2. (C) Representative confocal micrographs for each sample. Scale bars are 10µm. (D) Quantification of co-localization between LC3B and LAMP2; ****P(adjusted) < 0.0001, **P(adjusted) = 0.0014, ANOVA with Dunnett’s multiple comparisons test. (E) Immunoblots of Lyso-IP samples from sgNT or sgNPC1 293Ts treated with Torin1, rapamycin, or vehicle as in (C). (F and G) sgNT or sgNPC1 293Ts expressing control (shLuciferase) or Ragulator-specific (shLAMTOR5) shRNAs were fixed and stained with antibodies directed against LC3B and LAMP2. (F) Representative confocal micrographs for each sample. Scale bars are 10µm. (G) Quantification of co-localization between LC3B and LAMP2; ****P(adjusted) < 0.0001, ANOVA with Dunnett’s multiple comparisons test. (H) Immunoblots of Lyso-IP samples from sgNT or sgNPC1 293Ts treated with Torin1 or vehicle as in (C). (I and J) sgNT and sgNPC1 293Ts were pre-treated with Torin1 or DMSO for 24h before being treated with LLOMe (1mM) or vehicle (DMSO) for 10m. Cells were fixed and stained with antibodies directed against CHMP1A and TMEM192. (I) Representative confocal micrographs of LLOMe-treated cells. Scale bars are 10µm. (J) Quantification of co-localization between CHMP1A and TMEM192; ****P(adjusted) < 0.0001, ANOVA with Dunnett’s multiple comparisons test.

Due to the ability of mTORC1 to control lysosomal biogenesis and catabolism (Kim et al., 2011; Martina et al., 2012; Roczniak-Ferguson et al., 2012; Settembre et al., 2012), we next tested whether mTORC1 inhibition could alleviate at least some aspects of lysosomal dysfunction identified by our lysosomal proteomic analysis. This was clearly the case. Treating NPC1-null cells with Torin1 promoted clearance of autophagic material from the lysosomal lumen, as judged by both LC3B-LAMP2 double immunofluorescence (Figure 4C-4D and S5A-S5B) and by direct immunoblotting of immunopurified lysosomal samples from both NPC1-KO HEK-293T and MEFs (Figure 4E and S5C). Also, inhibiting mTORC1 signaling via shRNA-mediated knock down of LAMTOR5, which is essential for mTORC1 recruitment to and activation at the lysosome (but is not required for mTORC2-dependent signaling) (Anandapadamanaban et al., 2019; de Araujo et al., 2017; Bar-Peled et al., 2012; Rogala et al., 2019; Su et al., 2017), led to pronounced clearance of accumulated LC3 from the lumen of LAMP2-positive NPC lysosomes (Figure 4F-4G and S5D).

In contrast to mTORC1 inhibition via Torin1 or Lamtor5 knock down, little effect on lysosomal clearance was observed when cells were treated with the allosteric mTORC1 inhibitor, Rapamycin (Figure 4C-4E and S5A-S5C). Rapamycin incompletely inhibits mTORC1: whereas S6 kinase-dependent anabolic programs are efficiently suppressed by Rapamycin, protein synthesis triggered by mTORC1-dependent 4E-BP1 phosphorylation is largely unaffected, as is suppression of catabolic programs mediated by mTORC1-dependent phosphorylation of Unc1-like kinase (ULK1) and the master regulator of lysosomal biogenesis, transcription factor EB (TFEB) (Lawrence and Zoncu, 2019; Liu and Sabatini, 2020). The higher efficacy of Torin1 (and Lamtor5 depletion) over Rapamycin suggests that inhibition of protein synthesis and activation of the autophagy-lysosome system are both key for restoration of lysosomal proteolysis downstream of mTORC1 inhibition.

mTORC1 inhibition via Torin1 treatment also corrected the higher propensity for damage of NPC1 lysosomes, as shown by decreased recruitment of ESCRT III subunits both by immunoblotting of purified lysosomal samples and by double immunofluorescence for CHMP1A or CHMP1B and lysosomal markers (Figure 4H-4J and S5E-S5G). Given that mTORC1 inhibition reversed proteolytic failure but not cholesterol accumulation within NPC lysosomes, we conclude that the higher propensity of NPC lysosomes to undergo membrane damage primarily results from accumulation of undigested substrates within the lumen and not from altered fluidity of the membrane due to increased cholesterol content.

### 5. mTORC1 inhibition restores lysosomal function in iPSC-derived NPC neuronal cultures

Lysosomal dysfunction and its correction by mTORC1 inhibition were readily observed in actively proliferating cell lines. However, the cell type most compromised by loss of NPC1 are postmitotic neurons in the cerebellum, cerebral cortex and other brain regions (Walkley and Suzuki, 2004). To characterize the role of aberrant mTORC1 signaling on organelle homeostasis in a disease-relevant model of NPC, we deleted the NPC1 gene in induced pluripotent stem cells (iPSCs) using CRISPR/Cas9, and subsequently differentiated this population into neurons using a previously established method involving co-culture with a stromal cell line and FACS purification of neural populations (Ordonez et al., 2012) (Figure S6A-S6B). Neural stem cells and neurons derived by this differentiation method express neuronal lineage markers (nestin, MAP2, ßIII tubulin) (Figure S6B-S6C) and have been shown to be electrophysiologically active (Israel et al., 2012).

Consistent with the results in HEK-293T and MEFs and with previous reports (Elrick et al., 2012), iPSC-derived NPC neurons showed accumulation of undigested autophagic adaptors TAX1BP1 and GABARAP (Figure 5A-5D). Interestingly, unlike the HEK-293T model, iPSC-derived NPC neurons showed accumulation of autophagic adaptors both within and outside LAMP2-positive lysosomes, suggesting that defective autophagosome-lysosome fusion compounds with impaired lysosomal proteolysis in these cells (Sarkar et al., 2013) Overnight treatment of NPC neurons with Torin1 largely cleared intracellular TAX1BP1 and GABARAP aggregates, suggesting that mTORC1 inhibition is sufficient to restore the function of the autolysosomal system of neuronal cells (Figure 5A-5D).

**Figure 5.**
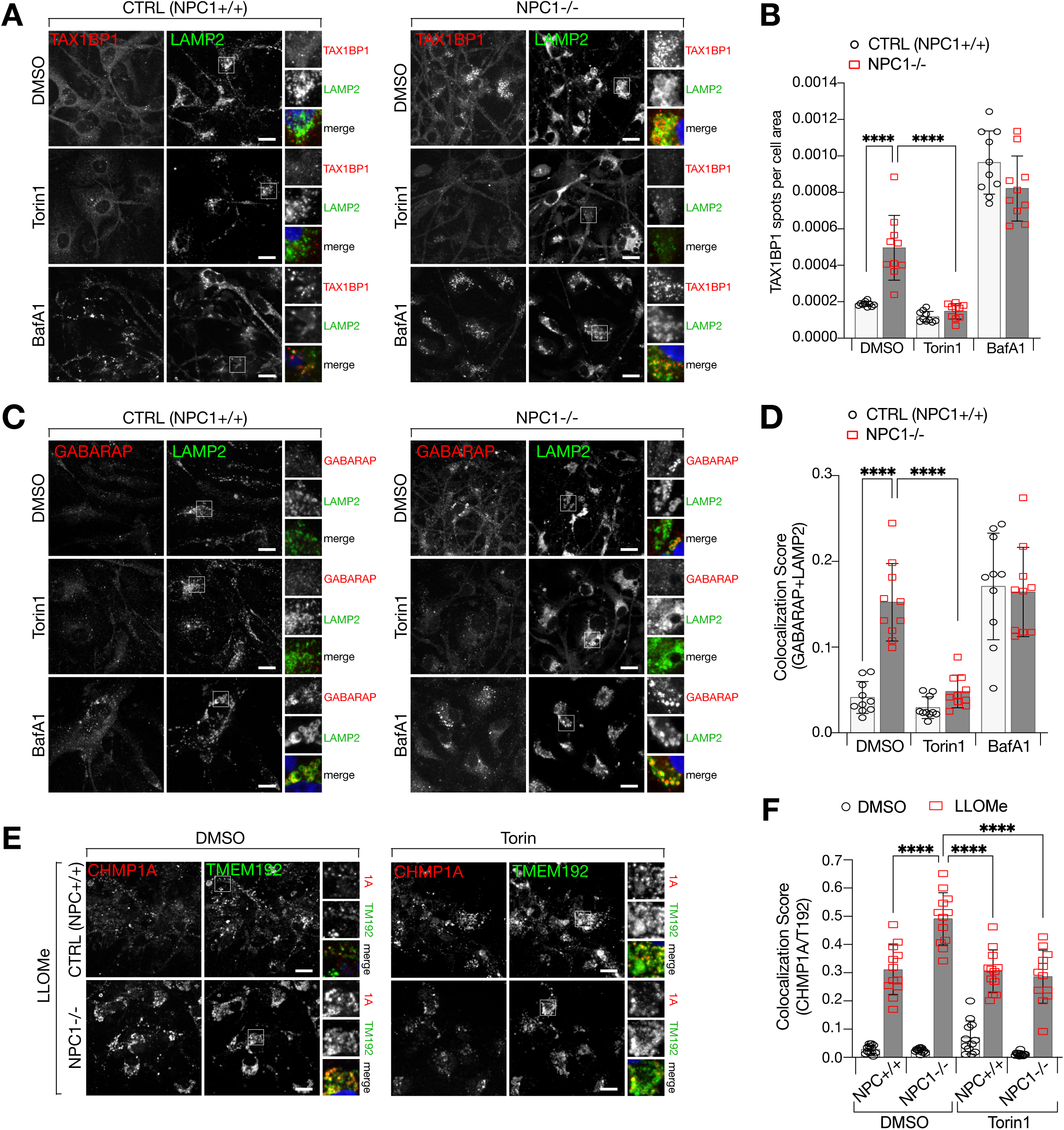
Inhibition of mTORC1 corrects lysosomal defects associated with loss of NPC1 in and iPSC-derived neuronal cell model. (A and B) Control or NPC1-/- iPSC-derived neural lineage cells were treated with Torin1 (250nM, 24h), BafA1 (500nM, 4h), or vehicle (DMSO) before being fixed and stained with antibodies directed against TAX1BP1, LAMP2, and MAP2 (not shown). (A) Representative confocal micrographs of each sample. (B) Quantification of the number of TAX1BP1 spots per cell area (defined by MAP2, see Figure S6C); ****P(adjusted) < 0.0001, ANOVA with Dunnett’s multiple comparisons test. (C and D) Control or NPC1-/- iPSC-derived neuronal lineage cells were treated as in (A) before being fixed and stained with antibodies directed against GABARAP and LAMP2. (C) Representative confocal micrographs of each sample. (D) Quantification of the co-localization between GABARAP and LAMP2; ****P(adjusted) < 0.0001, ANOVA with Dunnett’s multiple comparisons test. (E and F) Control or NPC1-/- iPSC-derived neuronal lineage cells were pre-treated with Torin1 or DMSO for 24h before being treated with LLOMe (1mM) or vehicle (DMSO) for 10m. Cells were fixed and stained with antibodies directed against CHMP1A and TMEM192. (E) Representative confocal micrographs of LLOMe- treated cells. (F) Quantification of co-localization between CHMP1A and TMEM192; ****P(adjusted) < 0.0001, ANOVA with Dunnett’s multiple comparisons test. Scale bars in all images are 20µm.

Similar to NPC1-depleted HEK-293T cells, iPSC-derived NPC neurons also had increased sensitivity to membrane damage, as shown by enhanced recruitment of ESCRT III to lysosomes upon treatment with LLOMe (Figure 5E-5F). As seen in non-neuronal lines, the hyper-sensitivity of iPSC-derived cells to lysosomal damage was also corrected by Torin1 treatment (Figure 5E-5F).

Thus, the compositional and functional defects revealed by our lysosomal proteomics in HEK-293T cells extend to NPC1-null neurons and are corrected by mTORC1 inhibition.

### 6. mTORC1 inhibition restores defective mitochondrial function in NPC1-null cells

Given the pervasive role of the autophagy-lysosome system in cellular quality control, the defective degradative capacity of NPC1-null lysosomes is expected to affect the integrity and function of other cellular compartments. Mitochondria are especially dependent on efficient lysosome-autophagy function for their morphological and functional homeostasis; accordingly, severe mitochondria defects have been reported in NPC1-null cell lines and in neuronal cultures derived from human embryonic stem cells (hESCs) (Ordonez et al., 2012; Yambire et al., 2019a). Moreover, dysregulated mTORC1 activity could contribute to mitochondrial dysfunction via increased translational burden, production of reactive intermediates and morphological alterations (Ebrahimi-Fakhari et al., 2016; Morita et al., 2013, 2017).

We analyzed our lysosomal proteomic datasets for the relative representation of mitochondrial proteins, defined by MitoCarta (Calvo et al., 2016) or Gene Ontology classification. By comparing their normalized peptide abundance, we discovered that mitochondrial proteins were significantly under-represented in NPC1-defective compared to control lysosomes (Figure 6A). In fact, when we ranked proteins based on their preferential enrichment in wild-type over NPC1 lysosomes, approximately 50% of the proteins in the top quartile were classified as mitochondrial, whereas only 23.6% of proteins in the other three quartiles were mitochondrial (Figure 6B). About half of the proteins classified as mitochondrial displayed the expected behavior of substrates, where their peptide count was higher in leupeptin:pepstatin than in DMSO (“mito substrates”, Figure 6A-6B). Thus, unlike other substrates that reach the lumen of NPC lysosomes but fail to be degraded, lysosomal delivery of mitochondria appears hampered in NPC, possibly reflecting disruption of the process of mitophagy (Ebrahimi-Fakhari et al., 2016; Khaminets et al., 2016; Ordonez, 2012).

**Figure 6.**
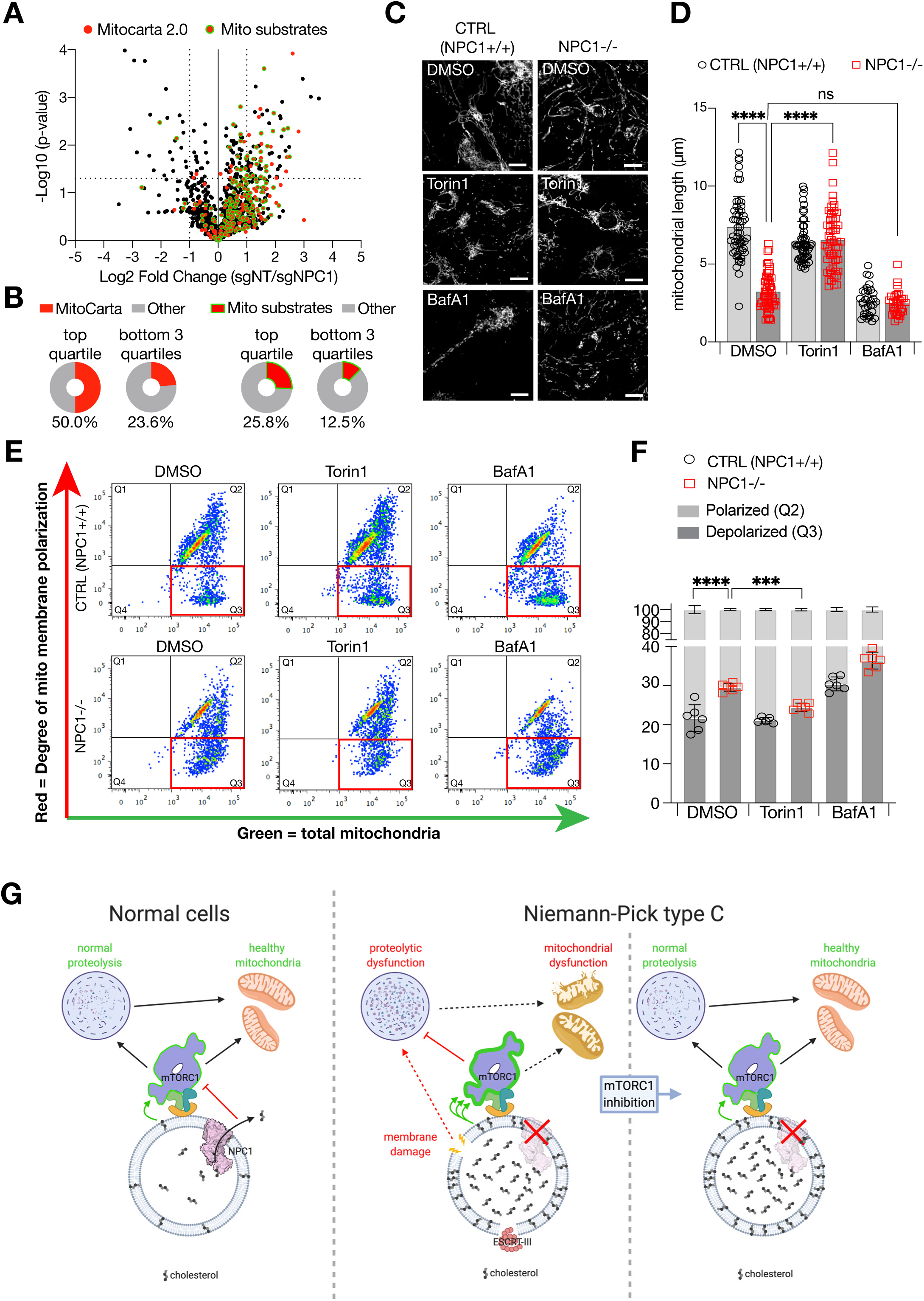
Mitochondrial morphology and function are disrupted by loss of NPC1 and is restored by inhibition of mTORC1. (A) Volcano plot of Lyso-IP proteomic data (from 1A-C) for the ratio of untreated (DMSO) sgNT/sgNPC1 LAMP1-normalized peptide counts. Proteins identified as mitochondrial, based on the human MitoCarta 2.0 database (Calvo et al., 2016), are shown as red circles. The subset of mitochondrial proteins that behave as substrates for lysosomal proteolysis are highlighted with a green outline. (B) Percentages of MitoCarta or “mito substrates” proteins that are in the top quartile (>75% enrichment) or remaining three quartiles (<75% enrichment) of proteins enriched in sgNT over sgNPC1 lysosomes. Total number of proteins is 1254, total number in top quartile is 62. (C and D) Control or NPC1-/- iPSC-derived neuronal lineage cells were treated with Torin1 (250nM, 24h), BafA1 (500nM, 4h), or vehicle (DMSO) before being fixed and stained with antibodies directed against TOM20. (A) Representative confocal micrographs of each sample. (B) Lengths of individual mitochondria were measured and quantified. (E and F) Control or NPC1-/- iPSC-derived neuronal lineage cells were treated with Torin1, BafA1, or vehicle as in (A) before being stained with the ratiometric mitochondrial membrane potential dye JC-10. (E) Dot plots showing fluorescence distribution of individual cells from one representative experiment. MMP-independent (“green”) fluorescence is shown on the x-axis and MMP-dependent (“red”) fluorescence is shown on the y-axis. (F) Percentages of cells classified as depolarized (Q3: green^high^, red^low^) or polarized (Q3: green^high^, red^high^). Values from individual replicates are shown as points, bars represent average values across all replicates. (G) Model illustrating the relationship between NPC1, lysosomal cholesterol, mTORC1 signaling and organelle homeostasis in both normal and NPC cells.

Consistent with defective mitophagy, the cytoplasm of iPSC-derived NPC neurons was disseminated with fragmented mitochondria, as shown by immunostaining with the mitochondrial marker Tom20 (Figure 6C-6D). Treatment with the v-ATPase inhibitor, BafA1, induced dramatic mitochondrial fragmentation in wild-type cells, consistent with an acute requirement for the lysosome in maintaining mitochondrial function (Ordonez et al., 2012; Weber et al., 2020; Yambire et al., 2019b). However, BafA1 only modestly increased the already high degree of mitochondrial fragmentation observed in NPC1-null cells, implicating lysosomal dysfunction as the main driver of this process (Figure 6C-6D). Further supporting this idea, overnight treatment with Torin1 restored mean mitochondrial length of NPC1-null neurons to wild-type levels, whereas it caused no change in mitochondria of their wild-type counterparts (Figure 6C-6D).

Similar to iPSC-derived NPC neurons, mitochondria of NPC1-defective HEK293T cells were highly fragmented, and their fragmentation was not further increased by BafA1-mediated v-ATPase inhibition (Figure S7A-S7B). Treating NPC1-defective HEK-293T cells with Torin1 re-established the tubular morphology of mitochondria (Figure S7A-S7B), whereas rapamycin was largely ineffective (Figure S7C-S7D).

Loss of mitochondrial integrity is often indicative of decreased mitochondrial membrane potential (MMP). Indeed, consistent with previous reports, mitochondria from NPC1-null iPSC-derived neurons showed significantly reduced MMP compared to wild-type cells, as determined by flow cytometry analysis of iPSC-derived neurons co-stained with the ratiometric MMP indicator, JC-10 (Figure 6E-6F). BafA1 treatment decreased the MMP of wild-type neurons, whereas it had a smaller effect on the MMP of NPC cells. Similar to mitochondrial fragmentation, Torin1 significantly rescued the MMP of NPC neurons, whereas it had a negligible effect on wild-type cultures. Thus, dysregulated mTORC1 signaling contributes to mitochondrial impairment in NPC neurons, and its pharmacological modulation is sufficient to restore both morphological and functional aspects of these organelles.

## DISCUSSION

Taken together, our results support a central role for the NPC1-cholesterol-mTORC1 signaling axis in the maintenance of organelle function and cellular homeostasis (Figure 6G). Export of lysosomal cholesterol by NPC1 is essential for optimal regulation of mTORC1 signaling outputs. In turn, mTORC1 signaling plays a critical role in sustaining the hydrolytic activities of the lysosome, the integrity of its limiting membranes, and the morphology and polarization of mitochondrial.

A key point that we address concerns the mechanisms via which NPC1 regulates mTORC1. We had previously shown that, in cells lacking NPC1, mTORC1 could not be switched off by cholesterol depletion, a phenotype consistent with both a cholesterol-transporting and a cholesterol-sensing role for NPC1 (Castellano et al., 2017; Lim et al., 2019). To determine which of these mechanisms is more likely to be correct, we systematically rescued NPC1-defective cells with different NPC1 mutants that lack cholesterol-exporting activities. These studies reveal a tight correlation between the cholesterol transporting function of NPC1 and its ability to regulate mTORC1 signaling, supporting a model in which NPC1 functions upstream of cholesterol, not downstream of it, in the mTORC1 pathway.

In turn, dysregulated mTORC1 signaling emerges as a key driver of organelle dysfunction downstream of NPC1 loss and the resulting lysosomal cholesterol buildup. This is shown by genetic and pharmacologic inhibition of mTORC1, which failed to correct cholesterol storage in the lysosomal lumen or on its limiting membrane but effectively corrected compositional and functional defects of the lysosome. How the lysosomal defects that we uncovered relate to each other will require further investigation. The depletion of several hydrolases (including proteases such as Cathepsin Z) provides a likely explanation for the profound proteolytic impairment of NPC lysosome revealed by accumulation of autophagic substrates. In turn, hydrolase depletion could stem from transcriptional inhibition or from their defective trafficking to the lysosome (Kobayashi et al., 1999; Saftig and Klumperman, 2009; Sleat et al., 2013). Alternatively, it is tempting to speculate that the increased propensity of NPC lysosomes to undergo membrane damage could result in leakage of lumenal contents, including resident hydrolases, a phenomenon suggested to occur in NPC as well as other physiological and pathological contexts (Chung et al., 2016; Hämälistö et al., 2020; Maejima et al., 2013; Sakamachi et al., 2017).

The primary factors driving membrane damage in NPC remain to be determined. In principle, alterations in membrane fluidity caused by the massive cholesterol accumulation on the limiting membrane of NPC lysosomes (as revealed by mCherry-D4H*) could increase the probability of membrane rupture, possibly through discontinuities between cholesterol-rich, crystalline-like microdomains and the surrounding membrane (Toulmay and Prinz, 2013; Tsuji et al., 2017). Damage could also result from undigested cargo accumulating within the lumen, a mechanism more similar to that induced by LLOMe, which assembles into membrane-piercing polymers within the acidic lysosomal lumen (Maejima et al., 2013; Radulovic et al., 2018; Skowyra et al., 2018). The observation that mTORC1 inhibition in NPC cells corrects lysosomal membrane damage and defective proteolysis without altering the cholesterol content of the lysosomal limiting membrane favors the latter possibility.

The observation that hydrolases mutated in other lysosomal storage disorders, such as acid lipase, beta-glucosidase and galactosidase are depleted from NPC lysosomes suggests the intriguing possibility that NPC compounds the pathogenic consequences of cholesterol storage with those of other diseases such as Wolman, Krabbe and Mucopolysaccharidosis III (Ballabio and Gieselmann, 2009), and could provide a mechanistic basis for the accumulation of neurofibrillary tangles, a pathogenic trait of Alzheimer Disease, in NPC brains (Walkley and Suzuki, 2004). More generally, hydrolase loss may be a common trait of several LSDs (Danyukova et al., 2018; Platt et al., 2018), a possibility that can be tested through the use of lysosomal profiling in cellular models of those diseases.

Given that mTORC1 inhibition decouples cholesterol storage from proteolytic failure and lysosomal membrane damage, a question that remains to be addressed is how NPC1-mTORC1 signaling controls these important functions of the lysosome. mTORC1 lies upstream of anabolic programs that could increase the proteolytic load of the lysosome, such as protein synthesis and ribosome biogenesis, while actively suppressing catabolic programs that could help restore lysosomal function, such as synthesis of lysosomal hydrolases and v-ATPase subunits by the MiT/TFE factors, TFEB and TFE3 (Düvel et al., 2010; Perera and Zoncu, 2016; Settembre et al., 2012). The failure of rapamycin, a partial mTORC1 inhibitor, at restoring lysosomal function in NPC cells suggests that simultaneous inhibition of biosynthetic programs and activation of catabolic ones downstream of mTORC1 may be required.

In our hands, overnight mTORC1 inhibition by Torin1 was sufficient to boost lysosomal proteolysis and correct susceptibility to damage. An interesting question is whether mTORC1 inhibition restores the function of pre-existing, cholesterol-filled lysosomes or rather promotes the formation of new ones. Answering this question, which may have implications for the use of mTORC1 inhibitors in clinical settings, will require follow-up analysis with single-organelle resolution.

Another important finding from this work concerns the role of mTORC1 signaling in maintaining mitochondrial morphology and function in NPC. Mitochondria are known to be severely impacted in many neurodegenerative diseases, and mitochondrial dysfunction is a major contributor to neuronal cell death observed in these contexts. However, there is limited understanding of the molecular processes that disrupt mitochondrial composition and function. mTORC1 stimulates translation of multiple nuclear-encoded mitochondrial transcripts (Morita et al., 2013), stimulates energy production as well as generation of anabolic intermediates in mitochondria (Ben-Sahra et al., 2016; Cunningham et al., 2007; Morita et al., 2013) and can indirectly promote mitochondrial fission (Morita et al., 2017). Moreover, by inhibiting autophagic initiation, elevated mTORC1 signaling can suppress the capture and clearance of damaged mitochondria, an observation in line with our lysosomal proteomic data (Ebrahimi-Fakhari et al., 2016; Perera and Zoncu, 2016). Thus, the ability of catalytic mTORC1 inhibitors to restore mitochondrial morphology and membrane potential likely stems from a combination of decreased translational burden and activity (associated with dampened production of reactive intermediates) and increased repair due to restoration of lysosomal proteolysis.

The restorative effects of mTORC1 inhibition on key functions in NPC cells suggests the attractive possibility that inhibiting this pathway could have therapeutic value. Our data clearly indicate that clinical derivatives of rapamycin (rapalogues) are unlikely to be beneficial, whereas the more potent and complete ATP-competitive inhibitors may be effective. However, the applicability of this class of inhibitors is limited by poor bioavailability and toxic off-target inhibition of mTORC2, which may lead to insulin resistance and other complications (Lamming et al., 2012; Liu and Sabatini, 2020). The recent development of new-generation, mTORC1-specific inhibitors with more complete inhibitory profiles than rapamycin (Chung et al., 2019; Mahoney et al., 2018; Rodrik-Outmezguine et al., 2016; Schreiber et al., 2019) may provide an avenue for safe and effective mTORC1 modulation in NPC and other lysosomal diseases.

## Acknowledgements

We thank all members of the Zoncu Lab for helpful insights. This work was supported by NIH R01GM127763 and R01GM130995 and a University of Notre Dame/APMRF grant to R.Z., a University of Notre Dame/APMRF 20174028 grant to M.P.O. and a 2019 AACR-Amgen Fellowship in Clinical/Translational Cancer Research (19- 40-11-SHIN) to H.R.S.

## Author Contributions

O.B.D, M.P.O., and R.Z. conceived of and designed experiments. O.B.D, H.R.S., C.Y.L., E.Y.W., M.K., C.F.M. and M.P.O. generated key reagents and performed the experiments. O.B.D, H.R.S., R.M.P., M.P.O. and R.Z. analyzed data and interpreted results. O.B.D., M.P.O and R.Z. wrote the manuscript.

All authors read and edited the manuscript.

## Methods Details

### Mammalian Cell Culture

HEK293T SGNT, HEK293T sgNPC1, MEF SGNT, and MEF NPC1-/- cells were maintained in DMEM (Gibco, 11995) supplemented with 10% (v/v) fetal bovine serum (Sigma, F0926) and 100 U/ml penicillin, and 100 µg/ml streptomycin (Gibco, 15140-122). All cells were cultured at 37°C in 5% CO2. Cells were free of mycoplasma and routinely tested using MycoAlert Mycoplasma Detection Kit (Lonza, LT07-318).

Drug treatments were performed as follows unless otherwise specified. Leupeptin (Alfa Aesar, J61188) and pepstatin A (MP Biologicals, 195368) were used at 20 µM each for 24 h. Bafilomycin A1 (Alfa Aesar, J61835) was used at 500 nM for 4 h. LLOMe (Sigma, L7393) was used at 1 mM for 10 m. Torin1 (Tocris, 4247) was used at 250 nM for 24 h, and rapamycin (Calbiochem, 553210) was used at 100 nM for 24 h.

### Cloning and Generation of Stable Cell Lines

A synthetic cDNA encoding TMEM192-RFP-3xHA (Lim et al., 2019) was cloned into the pLJM1 lentiviral vector. TMEM192-FLAG was cloned also cloned in the pLJM1 vector by amplifying the TMEM192 cDNA using primers to append a DYKDDDK (“FLAG”) peptide to the c-terminus of the resulting protein. Codon-optimized NPC1 cDNA containing a FLAG tag (Castellano, et al.) was subcloned into the pLVX lentiviral vector (Clontech). All NPC1 mutants were generated using the QuikChange II Site-Directed Mutagenesis Kit (Agilent, 200524).

Short-hairpin oligonucleotides (shRNAs) directed against LAMTOR5 (TRCN0000153443) or Luciferase (TRCN0000072243, used as a non-targeting control) were cloned into the pLKO.1 lentiviral vector (The RNAi Consortium, Broad Institute) according to the manufacturer’s instructions.

Expression of protein-encoding cDNAs or shRNA constructs was performed by stable lentiviral transduction. Lentivirus was generated by co-transfection of lentiviral vector carrying the construct of interest with lentiviral packaging plasmids (pMD2.G, Addgene 12259; and psPAX2, Addgene 12260) in a 5:3.75:1.25 ratio, respectively, using polyethylenimine (PEI). Viral supernatant was harvested 48 h after transfection, cleared by centrifugation, and concentrated using Lenti-X Concentrator (Clontech, 631231) according to the manufacturer’s instructions. Target cells were plated in 6-well plates in media supplemented with 8 µg/ml polybrene (Millipore, TR-1003-G) and concentrated virus. Virus-containing media was removed after 24 h and replaced with media containing 1.5 µg/ml puromycin. Protein expression or knockdown was confirmed by immunoblotting. For shRNA knockdown, cells were maintained in selective media for 3 days before use in assays in order to ensure complete knockdown.

### Lysosome Immunoprecipitation (Lyso-IP)

Lysosomes from cells expressing TMEM192-RFP-3xHA were purified as previously described (Lim et al., 2019). Briefly, cells were seeded in a 15cm at a density appropriate for them to reach confluency after 24h. All subsequent steps were performed on ice or at 4°C unless otherwise noted. Media was removed, cell monolayers were rinsed with ice-cold KPBS buffer (136 mM KCl, 10m M KH2PO4, pH 7.25, supplemented with Pierce Protease Inhibitor Tablets (Thermo, A32965)), scraped into 10 ml of KPBS and collected by centrifugation at 1500 rpm for 5 min. Pelleted cells were resuspended in a total volume of 1ml KPBS (supplemented with 3.6% (w/v) OptiPrep (Sigma, D1556)) and fractionated by passing through a 23G syringe 5 times followed by centrifugation at 2700 rpm for 10 min. Post-nuclear supernatant was harvested and incubated with 40 µl of anti-HA magnetic beads (Thermo, 88836, prewashed with KPBS buffer) with end-over-end rotation for 10 min. Lysosome-bound beads were washed two times with KPBS(+ OptiPrep) and two times with KPBS. For immunoblotting, samples were incubated with a 1:1 mixture of KPBS and 2x urea sample buffer (150 mM Tris, pH 6.5, 6 M urea, 6% SDS, 25% glycerol, 5% β-mercaptoethanol, 0.02% bromophenol blue) for 30 min at 37°C. For proteomics experiments, lysosomal immunoprecipitates were eluted from beads using 0.1% NP-40 in PBS for 30 min at 37°C, beads were removed and the resulting eluate was snap-frozen with LN2.

### Proteomics Analysis

For comparative analysis between treatment conditions and genotypes, minimum peptide abundance was set to 1 for all replicates. Experimental datasets were first compared to the proteomic dataset generated from anti-HA Lyso-IP performed on cells expressing TMEM192-FLAG (“blank” samples). Only proteins present with a combined average peptide abundance across both experimental samples >1.5-fold enrichment over blank samples were included in further analysis. Fold changes between experimental samples were then calculated, and the significance of these fold changes were calculated using a two-tailed unpaired t-test. For comparative analysis between genotypes, peptide counts from each replicate were additionally normalized to the peptide abundance of LAMP1 within each replicate, before calculation of fold changes and significance values. Data in all volcano plots are displayed as the log2 of the fold change, and the -log10 of the p-value.

The list of “cargo” proteins was generated by identifying all proteins whose abundance increased ≥2-fold upon inhibition of lysosomal proteolysis (leupeptin/pepstatin treatment). This list was cross-referenced to the dataset comparing leupeptin/pepstatin-treated to vehicle-treated sgNPC1 cells.

To analyze mitochondrial proteins present in Lyso-IP samples, the filtered datasets were cross-referenced with the Human MitoCarta 2.0 database (Calvo, et al., 2015; Pagliarini, et al., 2008). This list was further refined by eliminating proteins that did not obey expected behavior upon lysosomal proteolysis inhibition (i.e. any protein whose abundance decreased upon leupeptin/pepstatin treatment in either genotype was excluded) to generate the “mito substrates” subset. Quartile analysis is based on the enrichment of proteins in sgNT samples over sgNPC1 samples, where “top quartile” are proteins with an enrichment of >75% in sgNT samples, and “bottom three quartiles” are all other proteins. Percentages shown in pie charts represent the fraction of proteins identified in MitoCarta 2.0 database, or as “mito substrates”.

### Immunofluorescence

Cells were seeded on fibronectin-coated glass coverslips in 12-well plates at 150,000-300,000 cells per well, and allowed to attach overnight. Cells were treated with compounds at the specified concentrations and length of time as indicated before being fixed and stained. For LC3B, GABARAP, and TAX1BP1 staining cells were first fixed and permeabilized with ice-cold 100% methanol for 5 min at -20°C and then rinsed three times with PBS. For all other stainings, cells were first fixed with 4% paraformaldehyde (PFA) in PBS for 15 min at room temperature, rinsed three times with PBS, permeabilized with 0.1% (w/v) saponin in PBS for 10 min at room temperature, and rinsed three times with PBS. Primary antibodies were diluted into 5% normal donkey serum (Jackson ImmunoResearch, 017-000-121) and coverslips were labeled with this solution overnight at 4°C. Coverslips were rinsed three times with PBS and then labeled with fluorescently-conjugated secondary antibodies (diluted 1:400 in 5% normal donkey serum, PBS) for 45 min at room temperature, protected from light. Coverslips were rinsed with PBS six times (incubating in every other wash for 5 min at room temperature) and then mounted on glass slides using VECTASHIELD Antifade Mounting Medium with DAPI (Vector Laboratories, H-1200).

### Microscopy

All confocal microscopy was performed on a spinning-disk Nikon Ti-E inverted microscope (Nikon Instruments) system using a Plan Apo 60x oil objective. Images of fine cellular detail were acquired with an additional 1.5x magnifier. All images were acquired with an Andor Zyla-4.5 scientific complementary metal-oxide-semiconductor camera (Andor Technology) using iQ3 acquisition software (Andor Technology).

### Image analysis

For quantification of co-localization, 10-12 non-overlapping images were acquired from each coverslip. Raw, unprocessed images were imported into FIJI v.2.0.0-rc-69/1.52i and converted to 8-bit images, and images of individual channels were thresholded independently to exclude background and non-specific staining noise and converted to binary masks. Co-localization between lysosomes and the marker of interest was determined using the “AND” function of the image calculator. Data are plotted as the fraction of lysosomes that are positive for the marker of interest (the “Colocalization Score”).

For quantification of TAX1BP1 aggregates, thresholded images of the channel corresponding to MAP2 staining were used to generate a binary mask to define and measure the total area occupied by MAP2+ cells. Masks were then applied to independently thresholded images of channel corresponding to TAX1BP1 staining to exclude signal outside the defined cell area. Individual TAX1BP1 aggregates in the resulting image were counted using the “Analyze Particles” function, and data from individual frames are plotted as the average number of TAX1BP1 spots per MAP2+ cell area.

For mitochondrial perimeter measurements, individual cells were isolated into separate images and blinded before analysis. Each image was individually thresholded and converted to binary masks. The “Analyze Particles” function was used to identify and measure the perimeter of every particle in the resulting mask. Data are plotted as the average mitochondrial perimeter per cell analyzed. For mitochondrial length measurements, images were first blinded and then the length of individual mitochondria was measured manually. Data are plotted as the length of every individual mitochondria in each condition.

### Measurement of GALC activity

HEK293T cells were seeded at 10,000 cells per well in fibronectin-coated flat-bottom black 96-well plates with clear bottom (Greiner, 655090) and allowed to adhere overnight. Media was aspirated and then replaced with fresh complete growth media supplemented with 15 µM LysoLive GalGreen fluorogenic substrate (MarkerGene Technologies, M2776) and 50 nM LysoTracker Red DND-99 (Thermo Scientific, L7528) and incubated at 37°C for 2 h. Media was aspirated, wells were rinsed once with warm PBS, and then replaced with Imaging Buffer (136 mM NaCl, 2.5 mM KCl, 2 mM CaCl2, 1.3 mM MgCl2, 10 mM HEPES, pH 7.4). Endpoint fluorescence was measured on a Bio-Tek Synergy HT Multi-Mode Microplate Reader, using 485 nm/20 nm excitation with 528 nm/20 nm emission, and 570 nm/9 nm excitation with 590 nm/9 nm emission, read through the bottom of the plate. GalGreen fluorescence was normalized per well to LysoTracker Red fluorescence, and values from individual wells are plotted as points.

### Cholesterol starvation and replenishment

HEK293T cells were seeded in fibronectin coated culture dishes so they would reach 80-90% confluency at the start of the assay. For cholesterol depletion, cells were incubated in DMEM supplemented with 0.75% (w/v) methyl-β-cyclodextrin (MCD, Sigma C4555) and 0.5% (v/v) lipid-depleted serum (LDS) for 2 h. For cholesterol re-feeding cells were incubated with DMEM supplemented with 0.1% MCD and 0.5% LDS containing either 50 µM cholesterol (Sigma, C3045) or 100 mg/ml human LDL (Alfa Aesar, J65039), as indicated.

### Cell lysis and immunoblotting

Cells were incubated in lysis buffer (1% Triton X-100, 10 mM β-glycerol phosphate, 10 mM sodium pyrophosphate, 4 mM EDTA, 40 mM HEPES, pH 7.4, supplemented with Pierce protease inhibitor tablets) for 30 min at 4°C with rocking to ensure complete lysis. Lysates were harvested and cleared by centrifuging at 17,000g for 10 min at 4°C, protein concentration in the supernatant was measured by Bradford assay. Samples of equalized concentration were prepared for SDS-PAGE by addition of 2x Urea samples buffer or 5x sample buffer (235 mM Tris, pH 6.8, 10% SDS, 25% glycerol, 25% β-mercaptoethanol, 0.1% bromophenol blue). 5 µg of total protein from per sample was loaded per lane in a 12% Tris-Glycine gel (Thermo Scientific, XP00122) and resolved by electrophoresis in a Tris-Glycine running buffer (25 mM Tris, 190 mM glycine, 0.1% (w/v) SDS). For Lyso-IP samples 10% of the total immunoprecipitated material was loaded per lane, and 0.5% of total PNS was loaded per lane. Proteins were transferred to a PVDF membrane (Millipore IPVH00010), blocked with 5% non-fat milk in TBS-T, and incubated in primary antibodies (diluted in 5% milk in TBS-T) overnight at 4°C. Membranes were rinsed with TBS-T and incubated with horseradish peroxidase conjugated anti-rabbit or anti-mouse secondary antibodies (diluted in 5% milk in TBS-T) for 1 h at room temperature.

Membranes were washed again with TBS-T and incubated with Pierce ECL Western Blotting Substrate (Thermo Scientific, 32109) before being exposed to ProSignal ECL Blotting Film (Genesee Scientific, 30- 810L). For phosphorylation site specific antibodies, PBS-T was used in place of TBS-T for all steps, and antibodies were diluted in 5% BSA in PBS-T.

### Cholesterol labeling *in situ* with D4H*-mCherry and filipin

Recombinantly expressed GST-D4H*-mCherry was purified from BL21 *E. coli* as previously described (Lim et al., 2019). Labeling with D4H*-mCherry and filipin was also performed as previously detailed (Lim et al., 2019). Briefly, cells were plated on fibronectin-coated coverslips and treated as indicated before being fixed with 4% PFA in PBS for 15 min at room temperature. Coverslips were rinsed in PBS, and then selectively permeabilized by immersion in LN2 for 25 sec. Coverslips were then blocked with 1% BSA in PBS for 1 h at room temperature, followed by incubation in D4H*-mCherry, diluted 1:50 in 1% BSA/PBS, for 2 h at room temperature, protected from light. Coverslips were rinsed with PBS and then fixed again with 4% PFA/PBS for 10 min at room temperature. Coverslips were rinsed with PBS and filipin labeling was performed simultaneously with immunofluorescent staining of LAMP2. Primary and secondary antibodies were diluted in 1% BSA in PBS supplemented with 0.5 mg/ml filipin (Sigma, F9765) and performed each for 1 h at room temperature, rinsing the coverslips with PBS after each incubation. Coverslips were mounted on glass slides in VECTASHEILD Antifade Mounting Media (Vector Laboratories, H-1000).

### hIPSC generation and neuronal differentiation

Control hIPSC lines were derived from fibroblasts obtained from one healthy adult (J. Craig Venter) whose genome is fully sequenced and published. hIPSC lines are generated by four-factor reprogramming as previously described (Israel et al., 2012). Cell lines are examined for pluripotency by labeling with lineage specific markers Tra 1-81 (BD Bioscience), Oct-4 and Nanog (Santa Cruz). Pluripotency is assessed by embryoid body formation and staining for the 3 germ layers, endoderm (Alpha-fetoprotein, DAKO), mesoderm (smooth muscle actin, Millipore) and ectoderm (Nestin, Millipore). NSC and neurons were generated using previously described protocols and purified by FACS (Yuan et al., 2011). NSCs are stained for Nestin (Millipore) Sox1, Sox2 and Pax 6. Neurons are stained for MAP2 and TUJ. For each experiment involving hIPSC derived neurons, neurons from at least two independent differentiations were examined in duplicate or triplicate format.

### Generation of NPC1 knock-out hIPSCs

CRISPR/Cas9 gene editing was used to generate an NPC1 knock-out (KO) in the Craig Venter control hIPSC line. We inserted a frame-shift mutation in Exon 4 that engineered a premature stop codon leading to complete ablation of NPC1. Transfected hIPSCs were sorted based on GFP and Tra181 expression and sparsely plated onto 10cm MEF plates. Individual colonies were picked manually and transferred to 96-well plates. Candidates were screened by PCR and TOPO cloning, and positive hits were karyotyped to ensure genetic stability. Digital karyotype was normal. A microamplification on chromosome 1p was visually observed in parental CV line and NPC1 KO line but this was below threshold and considered to be an artifact. Ablation of NPC1 was confirmed by RTqPCR and Western Blot.

---

**Figure S1.**
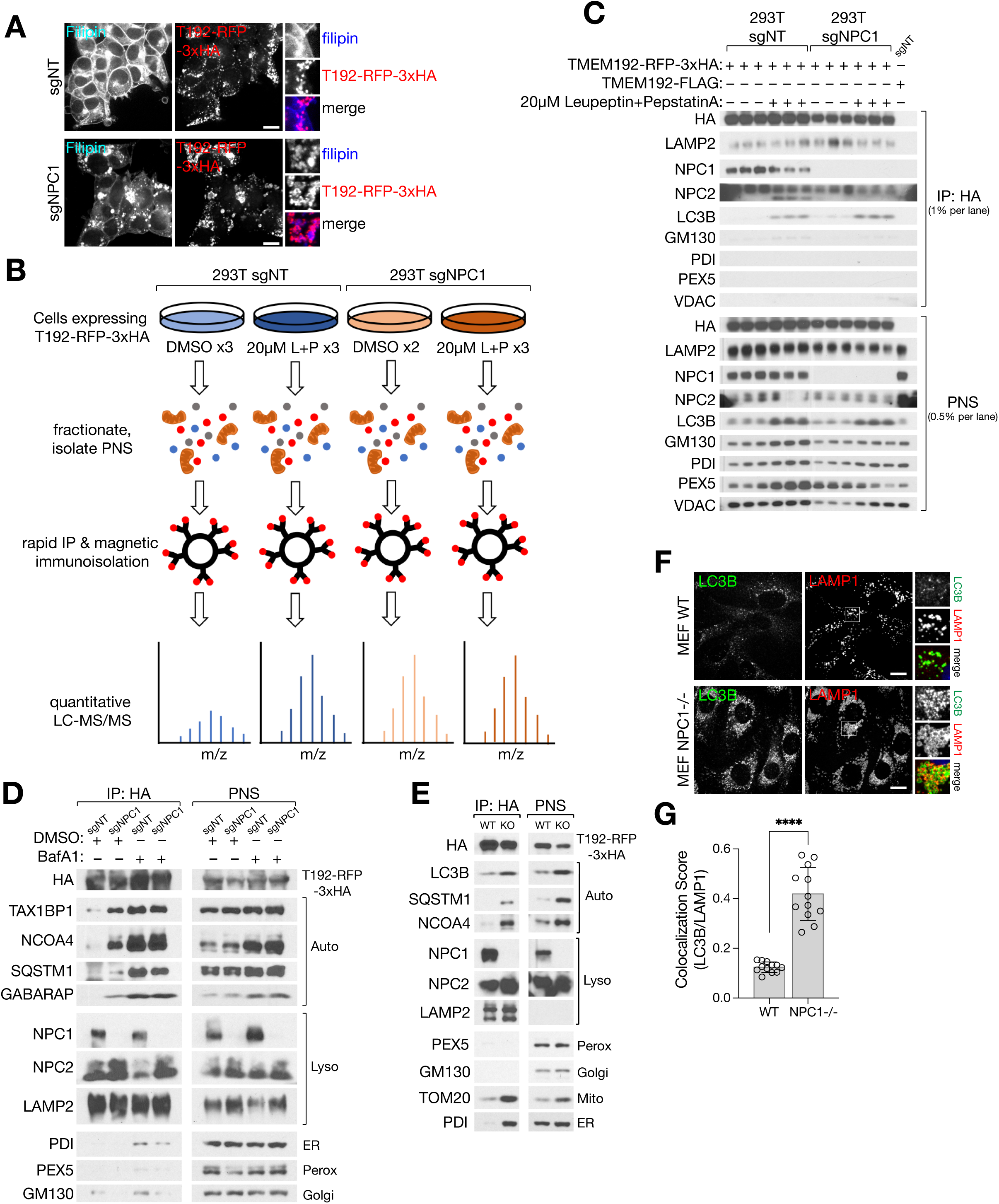
Loss of NPC1 results in a reduction of lysosomal proteolysis. Related to Figure 1. (A) Representative confocal micrographs from sgNT or sgNPC1 293Ts expressing TMEM192-RFP-3xHA and co-stained with filipin to reveal intracellular cholesterol distribution. (B) Schematic depicting protocol for proteomic analysis of lysosomal isolates generated with Lyso-IP. sgNT or sgNPC1 293Ts (expressing TMEM192-RFP-3xHA) were treated with leupeptin and pepstatin (L+P, 20µM each) or vehicle (DMSO) for 24h. Cells were mechanically fractionated and post-nuclear supernatant (PNS), containing lysosomes, was isolated by centrifugation. Intact lysosomes were purified by rapid immunoisolation, and lysosomal isolates were subjected to label-free quantitative proteomics. (C) Immunoblots of Lyso-IP samples used for label-free quantitative proteomics. Cells expressing TMEM192- FLAG were used as a negative control (only one replicate is shown). (D) Immunoblots of Lyso-IP samples from sgNT or sgNPC1 293Ts (expressing TMEM192-RFP-3xHA) treated with 500nM Bafilomycin A1 (BafA1) for 4h. (E) Immunoblots of Lyso-IP samples from sgNT or NPC1-/- (KO) MEFs (expressing TMEM192-RFP-3xHA). (F and G) WT and NPC1-/- MEFs were fixed and stained with antibodies directed against LC3B and LAMP1. (F) Representative confocal micrographs for each sample. (G) Quantification of co-localization between LC3B and LAMP1; ****P < 0.0001, two-tailed unpaired t-test with Welch’s correction. Scale bars in all images are 20µm.

**Figure S2.**
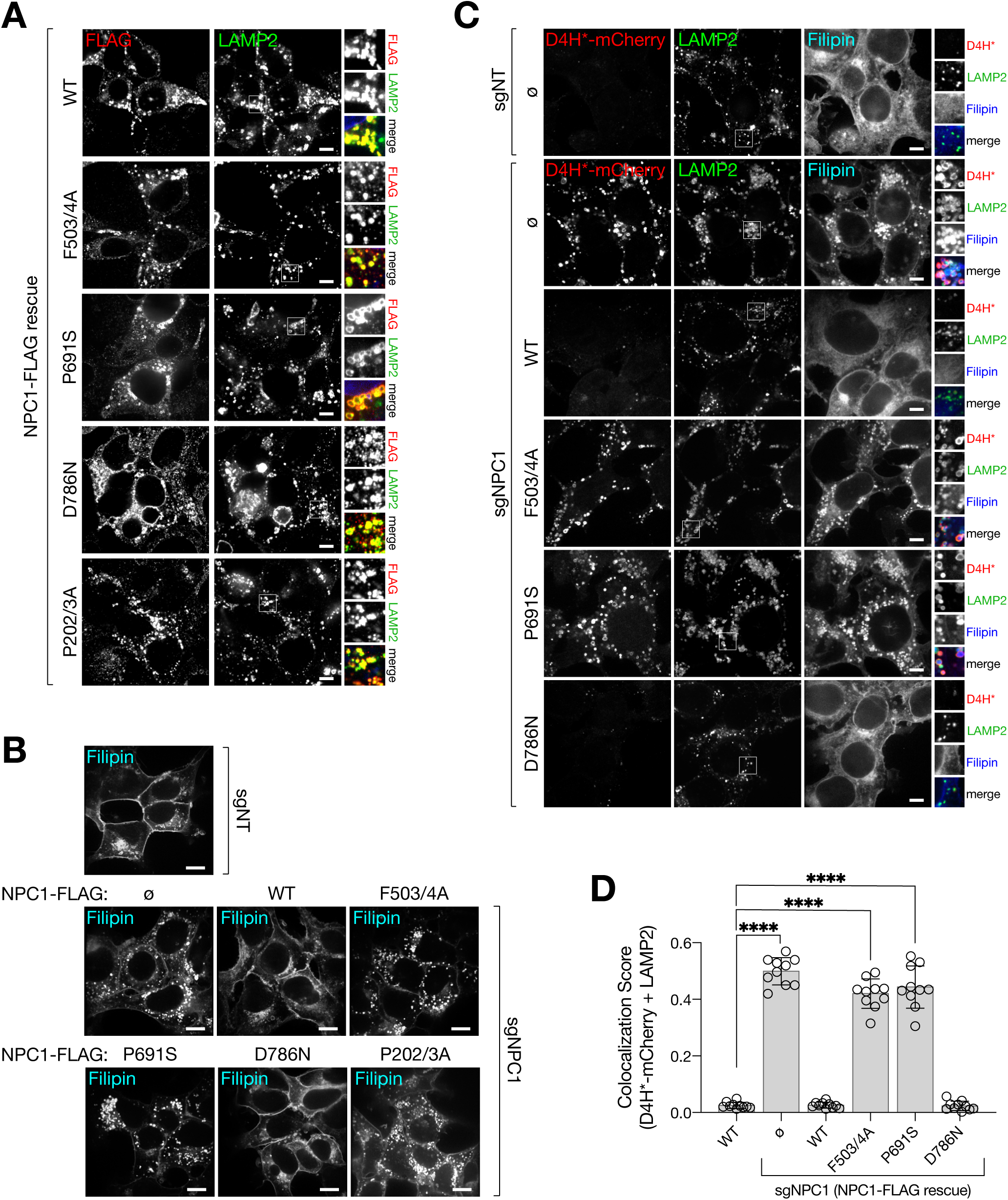
Reconstitution of NPC1-deficient cells with WT and mutant isoforms of NPC1 differentially regulates cholesterol export from lysosomes. Related to Figures 1 - 3. (A) sgNPC1 293Ts expressing the indicated NPC1-FLAG cDNA were fixed and stained with antibodies directed against FLAG and LAMP2. Scale bars are 10µm. (B) sgNT and sgNPC1 293Ts expressing the indicated NPC1-FLAG cDNA were fixed and cellular cholesterol deposits were stained with filipin. Scale bars are 20µm. (C and D) sgNT and sgNPC1 293Ts expressing the indicated NPC1-FLAG cDNA were fixed and semi-permeabilized with a liquid N2 pulse, followed by cholesterol labeling with D4H*-mCherry and filipin, and staining with antibodies directed against LAMP2. (C) Representative confocal micrographs for each sample. Scale bars are 10µm. (D) Quantification of the co-localization of D4H*-mCherry and LAMP2; ****P(adjusted) < 0.0001, ANOVA with Dunnett’s multiple comparisons test.

**Figure S3.**
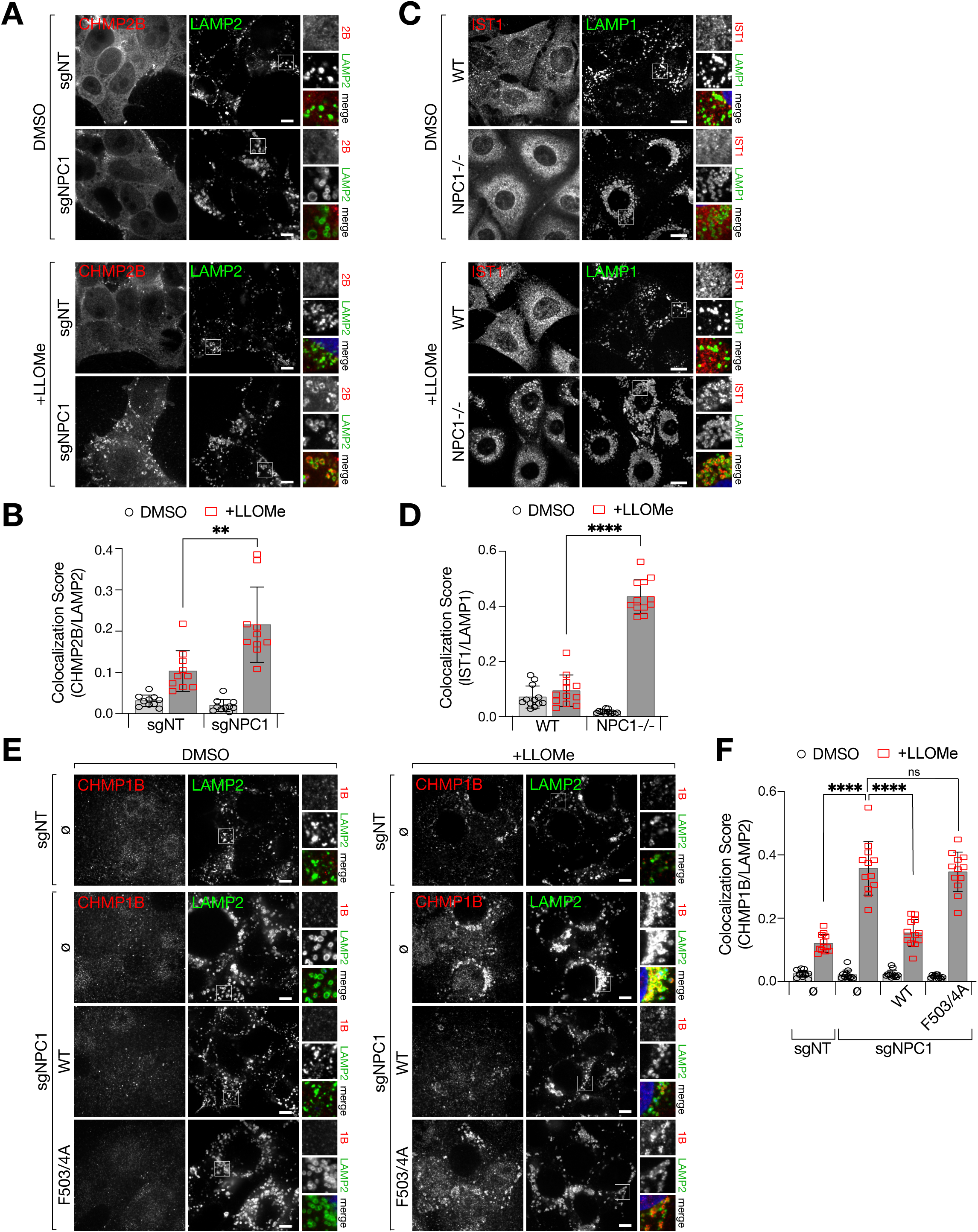
ESCRT-III recruitment to damaged lysosomes is enhanced by the loss of NPC1. Related to Figure 2. (A and B) sgNT or sgNPC1 293Ts were treated with 1mM LLOMe or vehicle (DMSO) for 10m before being fixed and stained with antibodies directed against CHMP2B and LAMP2. (A) Representative confocal micrographs for each sample. Scale bars are 10µm. (B) Quantification of the co-localization between CHMP2B and LAMP2; **P = 0.0043, two-tailed unpaired t-test with Welch’s correction. (C and D) WT and NPC1-/- MEFs were treated with 1mM LLOMe or vehicle (DMSO) for 10m before being fixed and stained with antibodies directed against IST1 and LAMP1. (C) Representative confocal micrographs for each sample. Scale bars are 20µm. (D) Quantification of the co-localization between IST1 and LAMP1; ****P < 0.0001, two-tailed unpaired t-test with Welch’s correction. (E and F) sgNT and sgNPC1 293Ts expressing the indicated NPC1-FLAG cDNA were fixed and stained with antibodies targeting CHMP1B and LAMP2. (E) Representative confocal micrographs for each sample. Scale bars are 10µm. (F) Quantification of the co-localization between CHMP1B and LAMP2; ****P(adjusted) < 0.0001, ANOVA with Dunnett’s multiple comparisons test.

**Figure S4.**
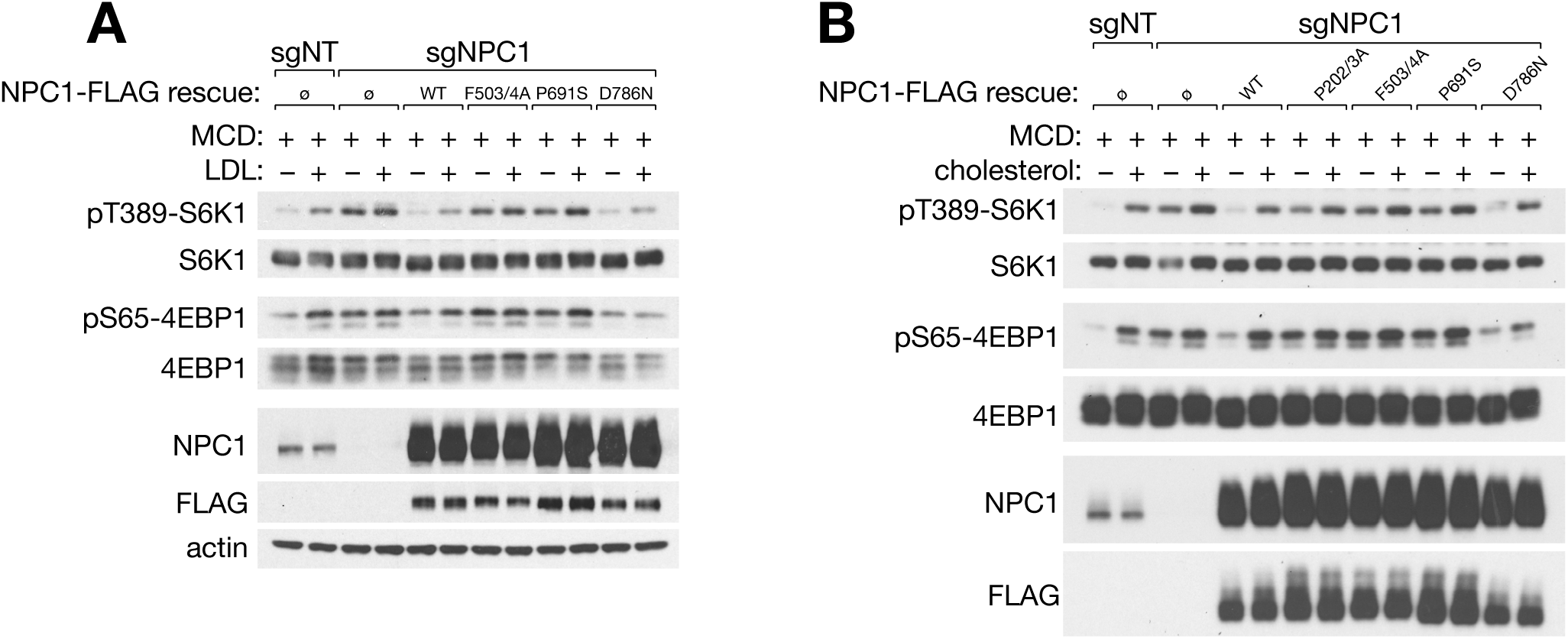
Re-expression of transport-defective NPC1 mutants fail to restore mTORC1 sensitivity to changes in cholesterol levels. Related to Figure 3. (A) Immunoblots from sgNT and sgNPC1 293Ts expressing the indicated NPC1-FLAG cDNA. Cells were depleted of sterols using methyl-β-cyclodextrin (MCD, 0.75% w/v) for 2h, followed by re-feeding for 2h with 0.1mg/ml human LDL particles in 0.1% MCD. (B) Immunoblots from sgNT and sgNPC1 293Ts expressing the indicated NPC1-FLAG cDNA. Cells were depleted of sterols using methyl-β-cyclodextrin (MCD, 0.75% w/v) for 2h, followed by re-feeding for 2h with 50µM cholesterol in complex with 0.1% MCD, as indicated.

**Figure S5.**
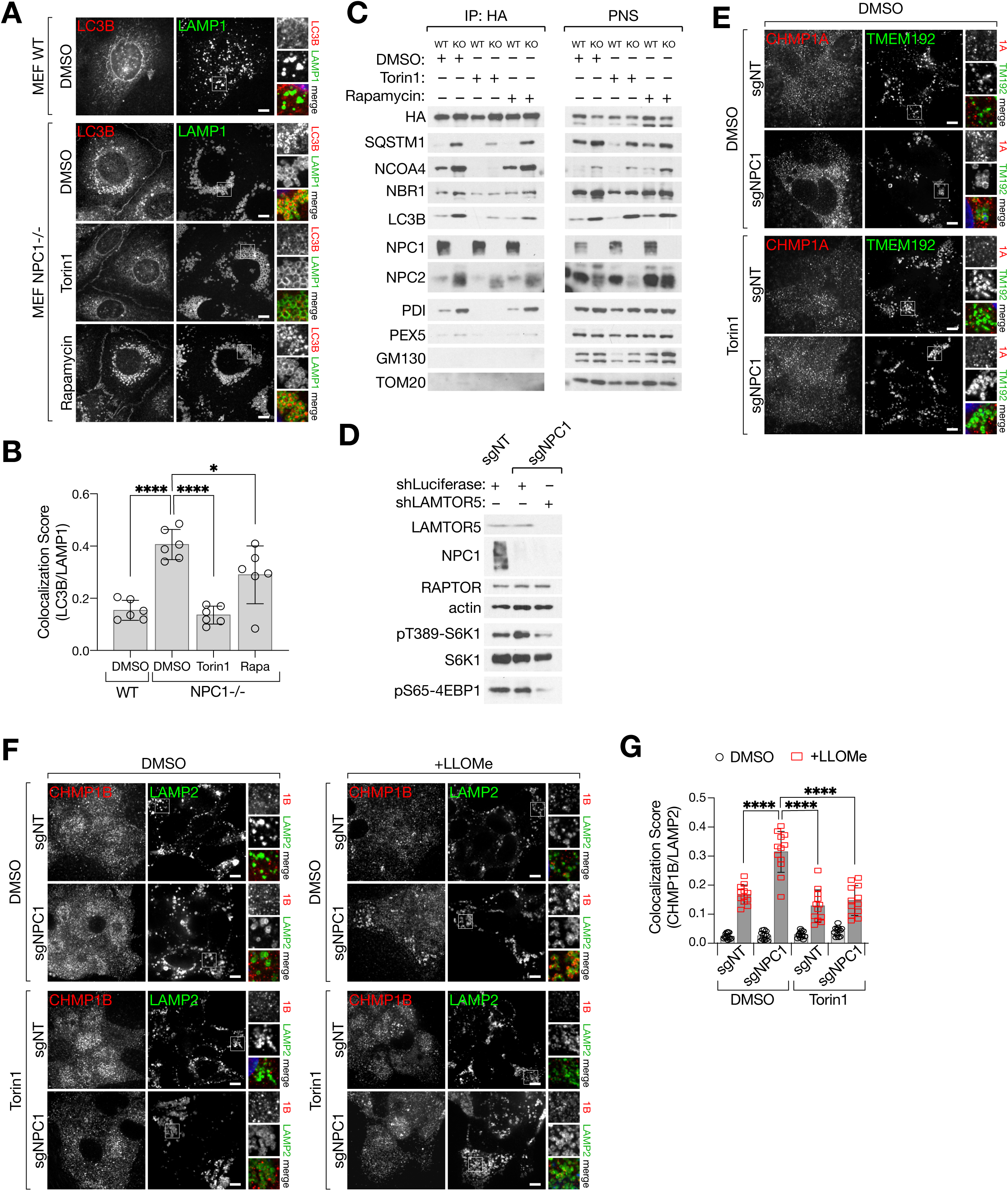
Effect of mTORC1 inhibitors on lysosomal proteolysis and membrane damage in WT and NPC1^-/-^ MEFs. Related to Fig. 4. (A and B) WT and NPC1-/- MEFs were treated with Torin1 (250nM, 24h), rapamycin (100nM, 24h), or vehicle (DMSO) as indicated before being fixed and stained for antibodies directed against LC3B and LAMP1. (A) Representative confocal micrographs for each sample. (B) Quantification of co-localization between LC3B and LAMP1; ****P(adjusted) < 0.0001, *P(adjusted) = 0.0426, ANOVA with Dunnett’s multiple comparisons test. (C) Immunoblots of Lyso-IP samples from WT or NPC1-/- (KO) MEFs treated with Torin1 (250nM, 24h), rapamycin (100nM, 24h), or vehicle (DMSO) as indicated. (D) Immunoblots of sgNT or sgNPC1 293Ts expressing control (shLuciferase) or Ragulator-specific (shLAMTOR5) shRNAs (E) Related to Figure 4I-J. Representative confocal micrographs of DMSO-treated sgNT or sgNPC1 293Ts. (F and G) sgNT and sgNPC1 293Ts were pre-treated with Torin1 or DMSO for 24h before being treated with LLOMe (1mM) or vehicle (DMSO) for 10m. Cells were fixed and stained with antibodies directed against CHMP1B and LAMP2. (F) Representative confocal micrographs of each sample. (G) Quantification of co- localization between CHMP1B and LAMP2; ****P(adjusted) < 0.0001, ANOVA with Dunnett’s multiple comparisons test. Scale bars in all images are 10µm.

**Figure S6.**
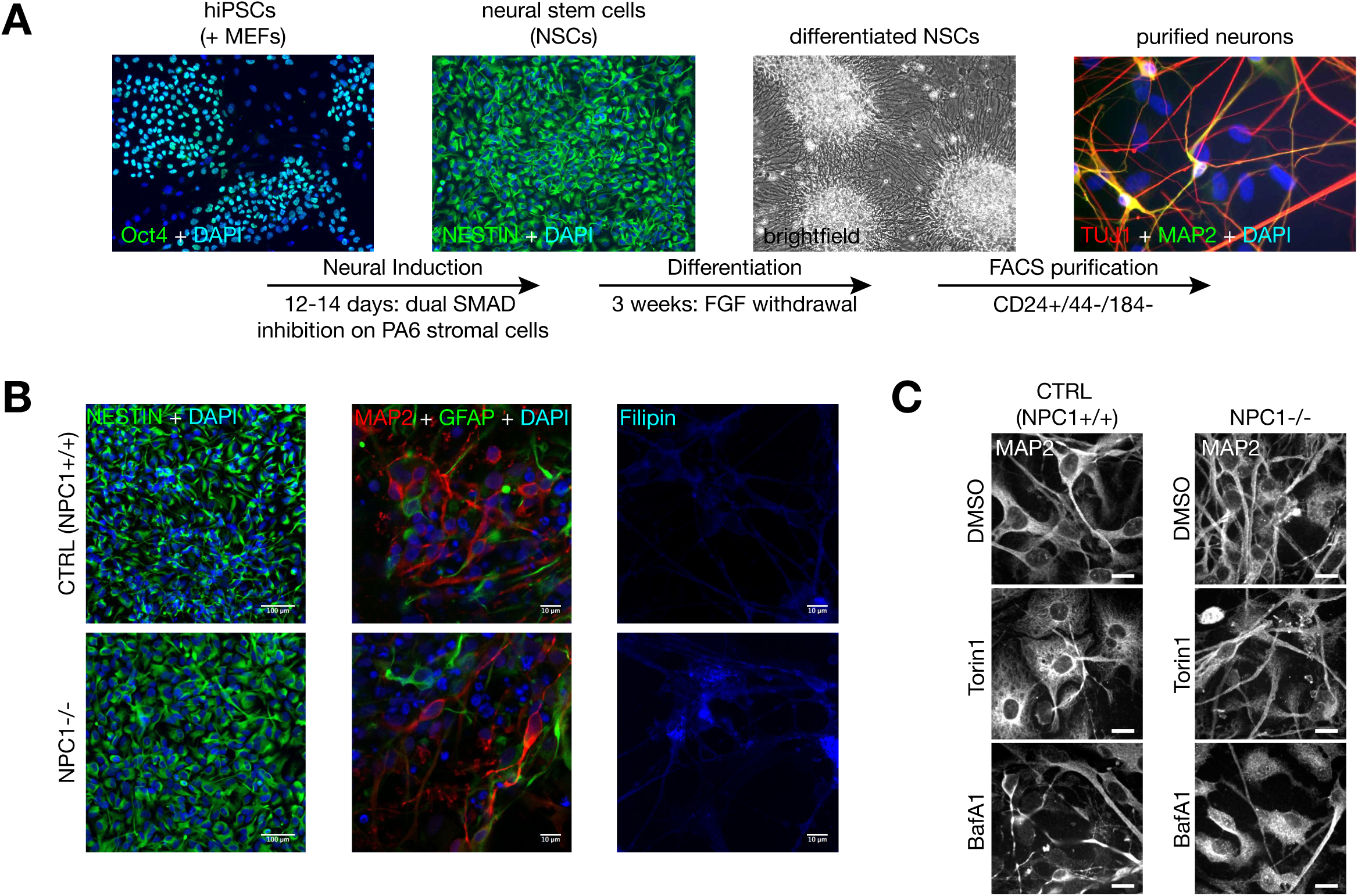
Validation of iPSC-derived neural lineage lines. Related to Figures 5 and 6. (A) Schematic depicting differentiation protocol for generating neural cell lineages from human iPSCs. Representative images depicting cellular phenotypes at each stage of differentiation are shown. (B) Validation of neural lineage differentiation for cell lines used in this study. NSCs were stained with Nestin, and differentiated neural cell populations were stained with MAP2 and GFAP. Cholesterol accumulation in NPC1-/- neural cells was validated by filipin staining. (C) Confocal images showing MAP2 staining corresponding to confocal images shown in Figure 5A. Scale bars are 20µm.

**Figure S7.**
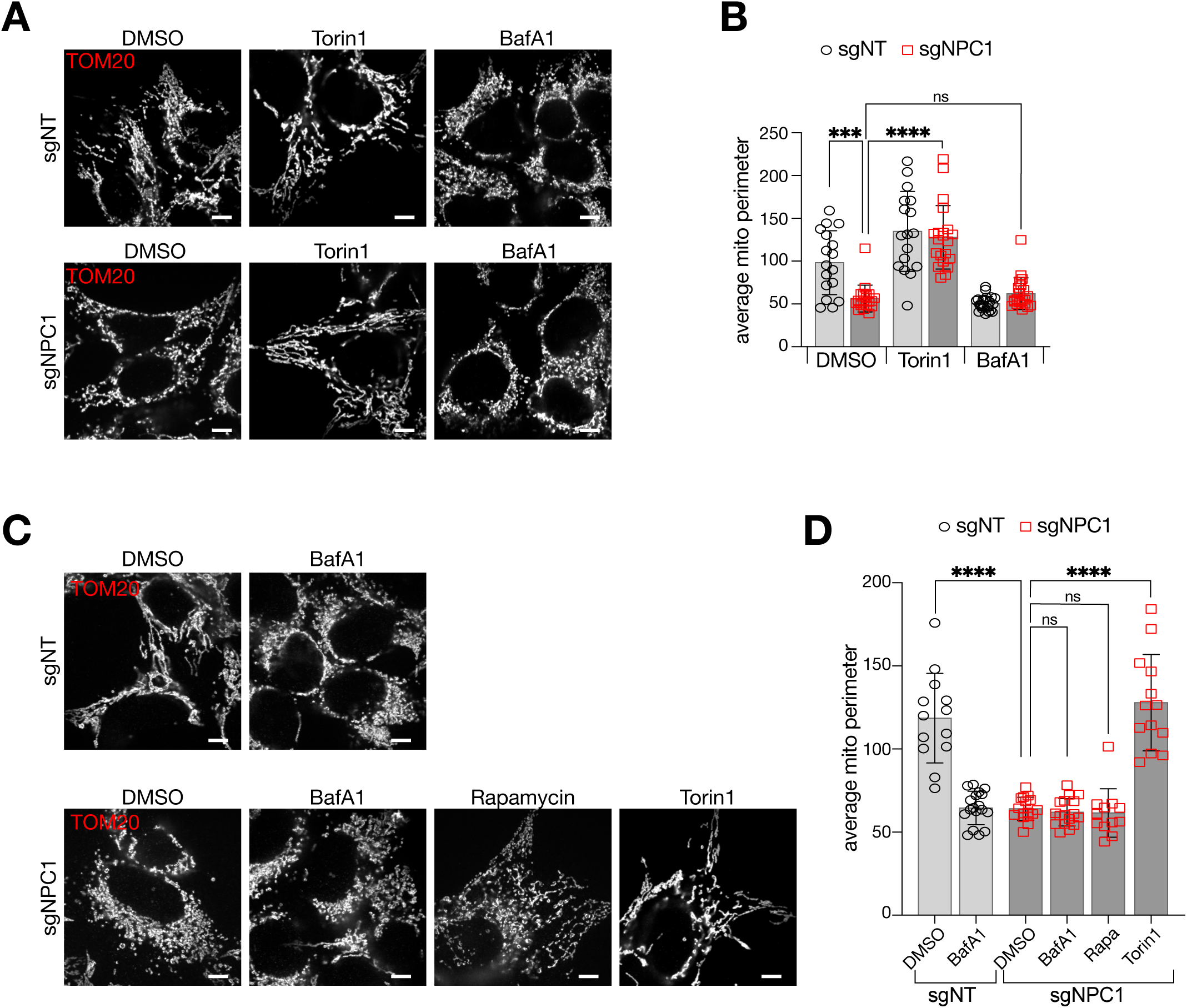
Mitochondrial morphology is perturbed in sgNPC1 cells and is restored by mTORC1 inhibition. Related to Figure 6. (A and B) sgNT and sgNPC1 293Ts were treated with Torin1 (250nM, 24h), BafA1 (500nM, 4h), or vehicle (DMSO) as indicated, before being fixed and stained with antibodies directed against TOM20. (A) representative confocal micrographs for each sample. (B) Quantification of average mitochondrial perimeter measured from TOM20 stained bodies. Number of cells in each condition are shown as individual points; ****P(adjusted) < 0.0001, ***P(adjusted) = 0.0002, ns = not significant, ANOVA with Dunnett’s multiple comparisons test. (C and D) sgNT and sgNPC1 293Ts were treated with BafA1 (500nM, 4h), Rapamycin (100nM, 24h), Torin1 (250nM, 24h) or vehicle (DMSO) as indicated, before being fixed and stained with antibodies directed against TOM20. (C) representative confocal micrographs for each sample. (D) Quantification of average mitochondrial perimeter measured from TOM20 stained bodies. Number of cells in each condition are shown as individual points; ****P(adjusted) < 0.0001, ns = not significant, ANOVA with Dunnett’s multiple comparisons test. Scale bars in all images are 10µm.

## Notes

### Competing Interest Statement

R.Z. is a scientific founder, shareholder and member of the scientific advisory board of Frontier Medicines, Corp.

